# Organyl 5′‑Phosphates in siRNA Guide Strands: Structure–Function Relationships Governing Anchoring in Argonaute 2 and Metabolic Stability

**DOI:** 10.64898/2026.02.13.705631

**Authors:** Theodore Carrigan-Broda, Luca F. R. Gebert, Samuel Hildebrand, Nozomi Yamada, Eric Luu, Jillian Caiazzi, Nicholas McHugh, Dimas Echeverria, Atish Wagh, Jonathan K. Watts, Anastasia Khvorova, Ian J. MacRae, Ken Yamada

## Abstract

Efficient siRNA loading into Argonaute2 (AGO2) requires a 5′-phosphate (5′-P) on the guide strand, yet this group is vulnerable to metabolic degradation in vivo. Although numerous chemical mimics of 5′-P have been reported, structural principles governing AGO2 interactions with organyl substituents on the 5′-P remain unclear. Moreover, structural determinants of 5′-P mimic recognition by known degradative enzymes (principally phosphatases and 5′-exonucleases) are also poorly understood. The 5′-P binding site of the AGO2 MID domain contains a stack of aromatic residues (Y527/F811/Y815), presenting a structural basis for augmenting canonical anchoring interactions. Herein, we systematically synthesized and characterized a diverse panel of organyl 5′-phosphates (5′-POR; R = 35 variable substituents) as guide strand 5′-P mimics designed to engage this unique hydrophobic pocket. Among the compounds evaluated, 5′-POR guide strands bearing methyl (Me) or phenylpropargyl (PhPrp) substituents are well-tolerated by AGO2 in cells. Previously uncharacterized 5′-P mimics, including 5′-phosphorothioate (5′-PS), phenylpropargyl 5′-phosphorothioate (5′-PS-PhPrp), and 5′-mesylphosphoramidate (5′-MsPA), maintain comparable AGO2 compatibility. All examined 5′-P mimics are markedly resistant to phosphatase, while 5′-POR variants and 5′-PS-PhPrp are also resistant to 5′-exonuclease degradation due to masking a negative charge of 5′-P. A crystal structure of a 5′-PO-PhPrp guide strand loaded into AGO2 reveals an unexpected network of π–π interactions between the rigid phenylpropargyl group and the targeted hydrophobic pocket of the MID domain. Collectively, these findings expand the functional chemical space of 5′-P mimics and define new modes for metabolically stabilizing the guide strand 5′-end while augmenting AGO2 MID anchoring.

**Graphical Abstract:** 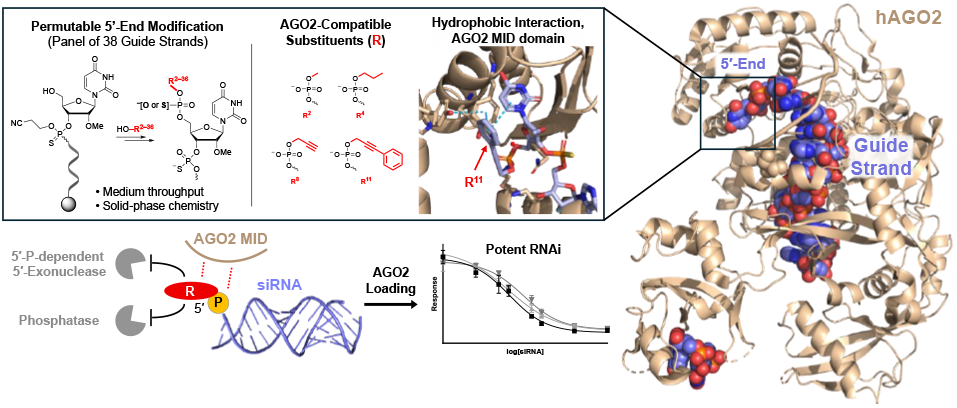

## INTRODUCTION

Short interfering RNAs (siRNAs) constitute a validated therapeutic modality, with eight approved drugs and many more in clinical development. ^1–5^ However, the multi-month pharmacological activity of siRNA drugs demands structural integrity of siRNA throughout intracellular trafficking, especially in the enzyme-laden endolysosomal compartment that sequesters then slowly releases endocytosed siRNA.^1,2^ Sustaining such metabolic stability in vivo entails chemical modification of all ribose residues (“full chemical modification,” typically 2′-*O*-methyl [2′-*O*-Me] or 2′-deoxy-2′-fluoro [2′-F] modifications) as well as terminal phosphorothioate (PS) internucleotide linkages;^1,6^ these modifications impart resistance to degradative nucleases while reducing immunogenicity.^2,6^ To effectuate any therapeutic gene silencing through the RNA interference (RNAi) pathway,^7–9^ the guide strand of siRNA must load into Argonaute2 (AGO2) to form the RNA-induced silencing complex (RISC),^1,10,11^ which specifically degrades complementary mRNA targets.^1,11^ Notably, RISC loading depends on a guide strand 5′-phosphate (5′-P; **Fig. 1A**),^1,9,12^ which anchors the guide strand in the AGO2 MID domain.^12,13^ However, the guide strand remains vulnerable to rapid phosphatase-mediated 5′-dephosphorylation.^2,14^ Conversely, kinase-mediated 5′-(re)phosphorylation of fully chemically modified guide strands is slow, so the dephosphorylated (RISC-incompatible) species predominates at steady-state.^2,12,15,16^ Additionally, slow but appreciable 5′-exonuclease activity can further erode guide strand integrity over an extended time.^17,18^ Thus, 5′-P mimics that preserve RISC compatibility while mitigating enzymatic degradation are advantageous, especially in extrahepatic tissues where siRNA accumulation is limited.^1,19^

**Figure 1.**
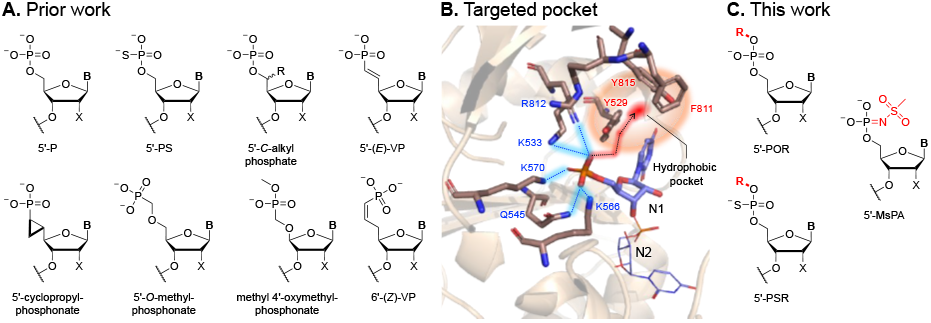
(**A**) Canonical phosphate (5′-P) and representative prior work of 5′-P mimics (**B** = nucleobase, X = 2′ modification). (**B**) Hypothesized mode of 5′-POR anchoring in AGO2 MID (adapted from PDBID: 4W5T). (**C**) This work: modular organyl 5′-phosphate (5′-POR), organyl 5′-phosphorothioate (5′-PSR) [R = variable organyl substituent], and 5′-mesylphosphoramidate (5′-MsPA).

A variety of 5′-P mimics, including 5′-*C*-alkyl phosphate and phosphatase-stable phosphonate derivatives, have been characterized for AGO2 loading (**Fig. 1A**).^20–22^ These studies collectively indicate that subtle structural alteration of 5′-P modulates interactions with AGO2 MID.^14,20^ Nevertheless, prior 5′-P mimic development mostly entailed modification of the phosphorus–ribose linkage; structural principles governing interactions of organyl substituents on 5′-P with AGO2 MID remain largely unexplored. In a prior study, masking a negative charge of the 5′-P with an aminoalkyl or photolabile organyl substituent was intended to transiently inactivate siRNA guide strands, and indeed markedly reduced RNAi activity.^23^ However, residual silencing activity was unexpectedly detected,^23^ suggesting that certain substituent architectures remain compatible with AGO2 loading. Likewise, the siRNA drug nedosiran features a 5′-P mimic with a methyl substituent (**Fig. 1A**) yet still exhibits high RNAi activity in vivo.^1,24^ These observations prompted our hypothesis that well-chosen substituents on the 5′-P could anchor in AGO2 MID with comparable efficiency to native 5′-P, warranting systematic exploration of the structural determinants of 5′-P mimic anchoring. Similarly, we previously showed that extended nucleic acid (exNA) modifications at the guide strand 3′-end are recognized by the AGO2 PAZ domain but not degradative 3′-exonucleases.^25^ We thus hypothesized that substituents on 5′-P may preserve AGO2 loading yet disrupt binding to phosphatase and 5′-exonuclease.

Crystallographic studies of RISC have revealed a conserved 5′-P binding pocket at the MID–PIWI interface, comprising cationic lysine residues (K533, K566, K570) and hydrogen bond donors (Y529, Q545, and ordered water molecules) that engage the non-bridging oxygens of 5′-P (**Fig. 1B**).^13^ Adjacent to this polar binding site lies an auxiliary hydrophobic region formed by stacked aromatic residues (Y529, F811, Y815),^13^ suggesting that select substituents appended to the 5′-P could interact favorably without disrupting canonical binding interactions. Hence, rational variation of 5′-P mimic structure constitutes a suitable platform to interrogate potential auxiliary anchoring modes within AGO2 MID.

In this study, we present a systematic structure–activity relationship (SAR) investigation of organyl substituents (R) on 5′-P (5′-POR), along with select 5′-phosphorothioate (5′-PS) and 5′-phosphoramidate (5′-PA) derivatives (**Fig. 1C**). By evaluating a panel of 35 diverse 5′-POR siRNAs, we demonstrated that AGO2 MID tolerates substantial stereoelectronic variability at the 5′-P, although some substituents substantially reduce RNAi potency. We also observed distinct patterns of susceptibility to phosphatase and 5′-exo-nuclease, governed differentially by charge valency and steric demand. Whereas resistance to 5′-exonucleases correlates principally with net charge of the 5′-P mimic, phosphatase susceptibility reflects both electrostatic and stringent steric constraints. Using atomic-resolution crystallographic analysis, we further show that the targeted hydrophobic pocket of AGO2 MID binds the rigid, aromatic phenylpropargyl (PhPrp) substituent through an unanticipated network of π–π interactions, explaining our observation of high RISC potency in cells. Additionally, AGO2 MID tolerates nonbridging sulfur or nitrogen atoms, as demonstrated by high RISC compatibility of 5′-PS, 5′-PS-PhPrp, and 5′-mesylphosphoramidate (5′-MsPA) guide strands. Finally, RISC pulldown experiments confirmed that a 5′-PO-PhPrp guide strand loads intact into RISC in cells. Collectively, these findings expand our understanding of the AGO2 MID interface with the guide strand 5′-end, define new principles for differentially modulating AGO2 anchoring and enzymatic susceptibility through rational chemical design, and extend the functional chemical space of 5′-P mimics.

## RESULTS AND DISCUSSION

### Synthesis of a 5′-POR guide strand panel

To characterize the effect of 5′-POR modification within a clinically relevant siRNA scaffold, guide strands were synthesized entirely from 2′-*O*-Me and 2′-F nucleotides with terminal PS linkages on UnyLinker support, conforming to a previously reported modification pattern (see supporting information).^26^ Guide strand sequences targeting either *HTT* or *MECP2* mRNAs were used.^26,27^ For 5′-POR modification of the full-length guide strand, we applied an “on-support phosphoramidite” strategy, wherein commercially available 2-cyanoethyl *N,N*-diisopropylchlorophosphoramidite reacts with the terminal 5′-hydroxyl (5′-OH) of the support-bound oligonucleotide to afford a universal 5′-phosphoramidite intermediate, which is subsequently coupled with diverse alcohols in medium throughput (**Fig. 2A**). Oligonucleotides were deprotected and cleaved from the solid support by the conventional ammonia-methylamine method. After filtering off the solid support and evaporating volatile bases, the obtained oligonucleotide was purified by ion-pair High-Performance Liquid Chromatography (HPLC), exchanged to the sodium salt via anion-exchange HPLC, and desalted (see supporting information). Purity and identity of all modified guide strands were validated via Liquid Chromatography–Mass Spectrometry (LC-MS; see **Table S11**).

**Figure 2.**
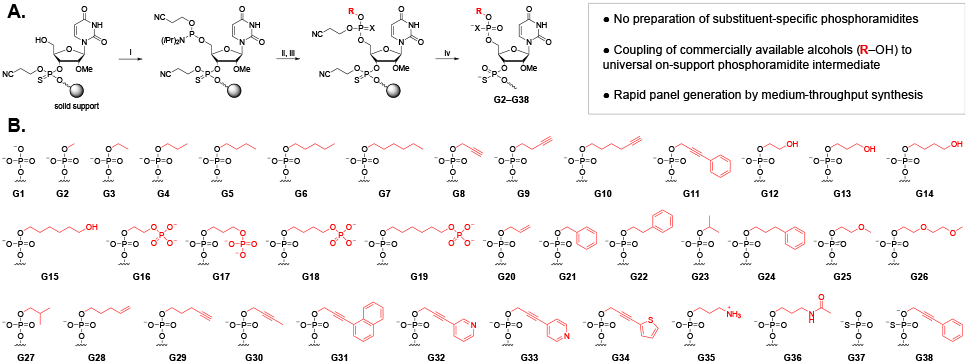
(**A**) Generalized on-support 5′-phosphoramidite scheme for 5′-POR and 5′-PSR modification of oligonucleotides (X = O or S): (i) 2-Cyanoethyl *N,N*-diisopropylchlorophosphoramidite, *i*Pr_2_EtN, MeCN, rt, 2h; (ii) ROH, 0.25 M ETT, MeCN, rt, 2h; (iii) oxidation:0.05 M I_2_ in pyridine-H_2_O (9:1, v/v), rt, 1 min; or sulfurization: 0.1 M DDTT in pyridine-MeCN (9:1, v/v), rt, 2.5 min; (iv) MeNH_2_-NH_4_OH (1:1, v/v), rt, 2 h. (**B**) 5′-end substructures of synthesized guide strands: 5′-P (**G1**), 5′-POR panel (**G2**–**G36**), 5′-PS (**G37**), and 5′-PS-PhPrp (**G38**) guide strands (**Table S11**).

### In cellula silencing activity of 5′-POR siRNA panel

Because guide strand anchoring in AGO2 governs all subsequent steps of the RNAi pathway—including RISC assembly and target mRNA binding and cleavage^28^—the silencing efficacy of 5′-POR siRNAs in cells serves as a direct indicator of their RISC compatibility. To evaluate silencing activity across our initial *HTT*-targeting 5′-POR guide strand panel (**G2**–**G23**; **Fig. 2B**), each guide strand was hybridized to a complementary passenger strand (**P1**) bearing a 3′-end cholesterol conjugate (**Table S1**), which facilitates passive uptake into cells.^26^ HeLa cells were treated with 5′-POR siR-NAs or positive control siRNA (5′-P, **G1**), and *HTT* mRNA expression in cellular lysate was quantified using the QuantiGene Singleplex system (ThermoFisher) and normalized to housekeeping gene (*HPRT*) expression.^26^ Although the silencing assay demonstrated appreciable RNAi activity for the entire panel (**Fig. 3A**), variability in efficacy by substituent suggests that AGO2 MID prefers certain 5′-POR structures. Using a target silencing threshold of >70%, we identified four hits from our guide strand panel, corresponding to methyl (Me; **G2**), propyl (Pr; **G4**), propargyl (Prp; **G8**), and phenylpropargyl (**G11**) substituents on the 5′-P (**Fig. 3C**). Although methyl and propyl are small alkyl substituents, propargyl and phenylpropargyl substituents contain a bulky, rigid alkyne moiety; the phenylpropargyl substituent also features extended π-conjugation with a phenyl ring.

**Figure 3.**
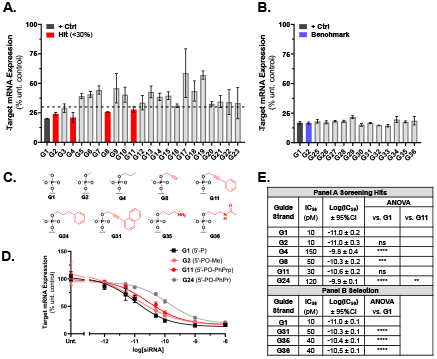
Silencing efficacy screening of (**A**) 5′-POR panel (**G2**– **G23**) and positive control (**G1**) [HeLa, 1.5 µM 3′-Chol siRNA^*HTT*^, 72 h]; (**B**) Panel addendum (**G25**–**G36**), positive control (**G1**), and **G2** benchmark [HeLa, 10 nM siRNA^*HTT*^, RNAiMax, 72 h]. Silencing efficacy = 100% – rel. target expression. (**C**) 5′-end substructures of hits from Panel A, **G11** analog (**G24**), and representative selection from Panel B. (**D**) Dose–response assay of hits (**G2, G11**), **G11** analog (**G24**), and control (**G1**) [HeLa, 0– 10 nM siRNA^*HTT*^, RNAiMAX, 72 h, *HPRT*-normalized]. (**E**) Potency and select pairwise IC_50_ comparisons from Panels A and B (one-way ANOVA, Šidák correction; α = 0.05; * p < 0.05, ** p < 0.01, *** p < 0.001, **** p < 0.0001).

Subsequently, these four hits and **G1** control were subjected to a dose–response assay to determine their cellular potencies. To further elucidate the SAR governing RISC loading, we also synthesized **G24** for comparison; this guide strand bears a phenylpropyl (PhPr) substituent (a flexible isostere of phenylpropargyl) on the 5′-P (**Fig. 3C**). Because steric bulk and lipophilicity of the substituents vary, all assessed siRNAs (**Table S2**) were lipofected into cells to equalize internalization; these siRNAs incorporated an unconjugated passenger strand (**P2**) to maximize lipofection efficiency. HeLa cells were treated with a seven-point dilution series of each siRNA, efficacy at each dose was determined as above, and potencies (half-maximal inhibitory concentration, IC_50_) were calculated from the semi-log dose–response curve (**Fig. 3D** and **Fig. S3**) and statistically compared (**Fig. 3E**). Our analysis indicates that introducing a methyl substituent to 5′-P does not appreciably affect silencing potency (**G1** vs **G2**), presumably because AGO2 MID readily accommodates this small steric change and hydrogen bonding between the alkylated oxygen and nearby cationic residues may compensate for the loss of the original electrostatic interaction. Interestingly, introduction of a phenylpropargyl substituent at the 5′-P also does not significantly affect potency (**G11** vs **G1**). Conversely, the siRNA with a phenylpropyl substituent exhibited markedly reduced potency (**G24** vs **G1**). The statistically significant fourfold difference between **G11** and **G24** potency indicates that the distinguishing stereoelectronic properties of phenylpropargyl—specifically its rigidity and aromatic π-conjugation rather than its size or lipophilicity—enhance RISC anchoring and underlie its unexpectedly high potency. Because neither **G2** nor **G11** differ significantly in potency from the **G1** control, their corresponding 5′-P mimics (5′-PO-Me and 5′-PO-PhPrp, respectively) were selected as lead 5′-POR variants for further characterization.

Considering this unexpected effect of π-conjugation on potency, we expanded the screening panel with twelve additional 5′-POR guide strands (**G25**–**G36, Fig. 2B**) to further explore SAR governing RISC compatibility; the substituents in this panel include numerous analogs of phenylpropargyl. Given the varied lipophilicity of these substituents, a 10 nM dose of each siRNA (**Table S3**) was lipofected into HeLa cells to equalize internalization; silencing efficacy was de-termined as above. All new test articles exhibited robust RNAi activity (**Fig. 3B**). To refine our SAR interpretation, we selected three of these siRNAs with representative guide strands (**G31, G35**, and **G36**; **Fig. 3C**) for a dose–response assay to determine silencing potencies as above (**Fig. S4**). The naphthylpropargyl substituent of **G31** extends the rigidity and π-conjugation of phenylpropargyl (**G11**). **G35** incorporates a previously reported “masked 5′-P” amino modification that retains partial efficacy in unmodified siRNA^23^ and can support post-synthetic *N*-acylation for the attachment of peptide or lipid conjugates. **G36** carries the corresponding acetylated derivative of **G35**. All selected guide strands exhibit modest but significantly reduced potency compared to **G1** control (**Fig. 3E**).

To confirm that 5′-POR siRNA potency is not sequence-dependent, lead 5′-POR guide strands (**G41**: 5′-PO-Me, **G42**: 5′-PO-PhPrp) and control (**G40**: 5′-P) targeting *MECP2* were synthesized and hybridized to an unconjugated passenger strand (**P3**). These siRNAs (**Table S5**) were subjected to dose–response assays as before (**Fig. S5**), which confirmed potencies were comparable to that of the 5′-P control.

### Synthesis and in cellula silencing potency of 5′-PS, 5′-PS-PhPrp, and 5′-MsPA siRNAs

The positive results from our 5′-POR panel screening prompted an expansion of the 5′-P mimic chemical space to further explore SAR governing RISC compatibility. For these additional 5′-P mimics, we used the same fully chemically modified guide strand as the 5′-POR panel (**Table S11**). First, we synthesized a 5′-PS guide strand (**G37, Fig. 2B**) as previously reported (**Scheme S1**)^29^ to benchmark the activity of other 5′-P mim-ics. Next, we extended our alkylation strategy to 5′-PS to assess the impact of the lead phenylpropargyl substituent. For the preparation of 5′-PS-PhPrp guide strand (**G38, Fig 2B**), the conventional oxidation step in the on-support phosphoramidite route was replaced with a standard sulfurizing step (**Fig. 2A** and **Scheme S2**). This sulfurization step is non-stereospecific and expected to produce a mixture of epimers.^30^ Likewise, the modification scheme for a 5′-MsPA guide strand (**G39**) was adapted from a previously reported internucleotide mesylphosphoramidate synthesis procedure.^31^ This strategy involves reaction of the 5′-hydroxyl with bis(2-cyanoethyl)-*N,N*-diisopropylphosphoramidite to afford an on-support 5′-phosphite intermediate, which then yields the protected 5′-MsPA product via an oxidative Staudinger reaction with mesyl azide (**Scheme S6**). 5′-MsPA provides particularly informative insights into RISC compatibility due to its distinctive stereoelectronic features, including a mesyl substituent on a non-bridging nitrogen rather than oxygen, a net charge of −2 (like canonical phosphate), and isosterism with pyrophosphate.

Potency of siRNA (**Table S4**) with **G37, G38**, or **G39** was evaluated in cellula via dose–response assay (**Fig. S6**). 5′-PS exhibited potency comparable to 5′-P (**G1** vs **G37**), indicating that the increased radius and polarizability of its nonbridging sulfur do not perturb RISC anchoring. Similarly, 5′-PS-PhPrp displayed potency comparable to its 5′-PO-PhPrp analog (**G11** vs **G38**), although the assay does not distinguish relative activities of the two presumed 5′-PS-PhPrp epimers. Despite its large steric demand, 5′-MsPA was also comparably potent to 5′-P (**G1** vs **G39**). Collectively, these results demonstrate that AGO2 can accommodate 5′-P mimics with markedly diverse stereoelectronic profiles.

### In vitro enzyme susceptibility of 5′-P mimics

Because metabolic stability is a principal determinant of successful siRNA drug development,^19^ 5′-P mimics that protect the guide strand 5′-end from both phosphatases and 5′-exonucleases are clinically relevant. Most previously reported 5′-P mimics (like 5′-*E*-VP) were designed to resist phosphatase through non-scissile covalent linkages (namely phosphorus–carbon bonds);^20^ inhibiting 5′-P mimic binding to phosphatase and 5′-exonuclease would also confer metabolic stability, but prior studies did not emphasize this approach. Another study found that 5′-*E*-VP confers siRNA guide strands with modest protection from 5′-exonuclease in vitro.^17^ However, the impact of other 5′-P mimics on 5′-exonuclease-mediated oligonucleotide degradation remains uncharacterized, despite pertinence to both siRNA and antisense oligonucleotide (ASO) drug development.^19,32^

To probe the SAR governing phosphatase susceptibility, lead 5′-POR (**G2**: 5′-PO-Me, **G11**: 5′-PO-PhPrp) and control (**G1**: 5′-P) guide strands (**Table S3**) were incubated with a bovine alkaline phosphatase (Quick CIP, New England Biolabs) at 37 °C. Samples were collected at five timepoints over 1 h and quenched. Intact 5′-POR or 5′-P guide strand and 5′-OH degradant within samples were identified by retention time on analytical HPLC by comparison to relevant standards, then quantified by *A*_260_ and plotted. 5′-PO-Me (**G2**) and 5′-PO-PhPrp (**G11**) guide strands exhibited no detectable degradation, whereas 5′-P (**G1**) was completely dephosphorylated to 5′-OH within 1 h under these conditions (**Fig. S7**). Given marked steric differences among the substituents tested, our results suggest that any substituent is sufficient to abrogate phosphatase-mediated degradation. This observation is consistent with prior reports, as typical nucleic acid phosphatases (phosphomonoesterases) display limited activity against phosphodiester substrates.^33,34^

To assess phosphatase susceptibility over a longer time course, lead 5′-POR (**G2**: 5′-PO-Me, **G11**: 5′-PO-MePhPrp), 5′-PS (**G37**), 5′-PS-PhPrp (**G38**), and 5′-MsPA (**G39**) guide strands (**Table S4**) were incubated with the same phosphatase at 37 °C for 12 h, then quenched, while the control (**G1**: 5′-P) was incubated for 1 h. These samples were characterized via LC-MS analysis and compared to the initial time point. Consistent with prior results, the 5′-P control (**G1**) was completely dephosphorylated to 5′-OH within 1 h, whereas all other 5′-P mimics exhibited no detectable degradation over 12 h (**Fig. 4A**). These results indicate that the phosphatase is highly selective for its native 5′-P substrate, with even 5′-PO-Me and 5′-PS guide strands remaining intact under the conditions tested. The observed selectivity is consistent with sensitivity to steric constraints or charge distribution.

**Figure 4.**
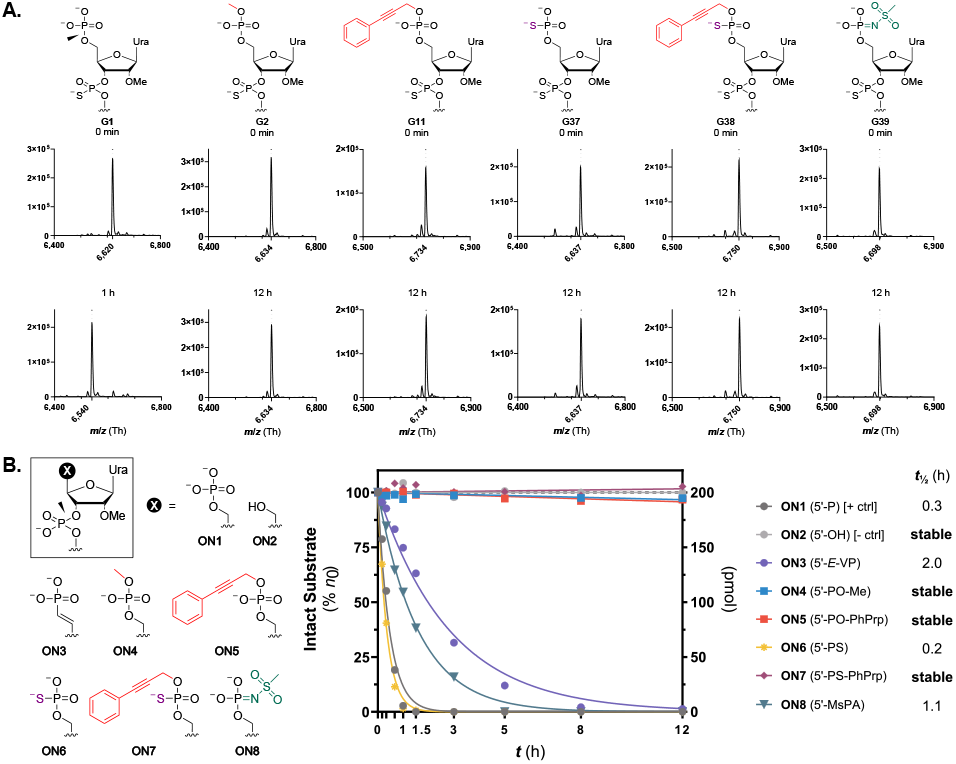
(**A**) Phosphatase susceptibility of lead 5′-POR, 5′-PS, 5′-PS-PhPrp, and 5′-MsPA guide strands over 12 h and 5′-P control over 1 h (0.1 mU/mL QuickCIP, LC-MS analysis). (**B**) Exonuclease susceptibility of lead 5′-POR, 5′-PS, 5′-PS-PhPrp, 5′-MsPA, 5′-P positive control, and 5′-OH negative control oligonucleotides (no PS linkages; 30 U/mL XRN1, HPLC analysis). Scissile bond indicated with arrowhead.

To probe the SAR of 5′-exonuclease susceptibility, 5′-*E*-VP (**ON3**), lead 5′-POR (**ON4**: 5′-PO-Me, **ON5**: 5′-PO-PhPrp), 5′-PS (**ON6**), 5′-PS-PhPrp (**ON7**), 5′-MsPA (**ON8**), positive control (**ON1**: 5′-P), and negative control (**ON2**: 5′-OH) fully oligonucleotides without terminal PS internucleotide linkages (**Table S5**) were synthesized and incubated with XRN1 (New England Biolabs), the principal cytosolic 5′-phosphate–dependent 5′-to-3′ exoribonuclease.^35^ Samples were taken at ten timepoints and quenched, then intact oligonucleotide was quantified via analytical HPLC (**Fig. 4B**). Intriguingly, only the oligonucleotides bearing a 5′-P mimic with an organyl substituent, namely 5′-PO-Me (**ON4**), 5′-PO-PhPrp (**ON5**), and 5′-PS-PhPrp (**ON7**), were resistant to XRN1 degradation. Conversely, all tested oligonucleotides with a dianionic 5′-P mimic, including 5′-*E*-VP (**ON3**), 5′-PS (**ON6**), 5′-MsPA (**ON8**), and 5′-P control (**ON1**), were rapidly and completely degraded by XRN1 (*t*_½_ < 2h) under the conditions tested.

These findings suggest that masking one of the negative charges of 5′-P with a substituent prevents degradation by XRN1, whereas mere steric bulk without a masked charge (as with 5′-MsPA) is not inhibitory. Based on a prior crystal structure of XRN1 with a 5′-P DNA substrate (**Fig. S8A**),^35^ we conjecture that the 5′-P binding pocket can accommodate a gauche substituent on 5′-P, but the masked charge reduces electrostatic attraction to K93, R100, and R101, resulting in shallower binding. This displacement may misalign the π–π stack formed by H41 and the substrate 5′-end nucleobases; alternatively, the π–π stack may rotate like a hinge to accommodate the displacement (**Fig. S8B**). These spatial changes likely perturb crucial interactions between the first internucleotide linkage of the substrate and a bound magnesium in the XRN1 active site, markedly reducing catalysis or processivity.

### In cellula metabolism and RISC loading of 5′-PO-PhPrp siRNA

Although the potent cellular RNAi activity of our lead 5′-POR siRNAs indirectly evinces RISC compatibility, these results do not exclude the possibility of intracellular conversion to 5′-P through hydrolysis of the substituent. To determine whether a 5′-POR guide strand retains its substituent after RISC loading, cells were treated with an siRNA (**D28**; see **Table S2**) with a 5′-PO-PhPrp guide strand (**G11**), then RISC was captured using a validated AGO2 pulldown approach, isolated, and analyzed by LC-MS (**Fig. 5A**).^36^ Briefly, HeLa cells were lipofected with the siRNA, incubated for 48 h, and lysed under non-denaturing buffer conditions. The total RISC pool was captured from the lysate (input) on beads via an Argonaute-binding fusion protein (GST-T6B), and the beads were washed to remove unbound matrix (supernatant), including unloaded guide strands. To corroborate RISC capture efficiency, an aliquot of the bead adsorbate (pulldown fraction) was analyzed with the input and supernatant via Western blot (**Fig. S9**), confirming AGO2 depletion in the supernatant and enrichment in the pulldown fraction. Likewise, unbound protein depletion in the pulldown fraction was corroborated via a control protein (GAPDH) in the blot.

**Figure 5.**
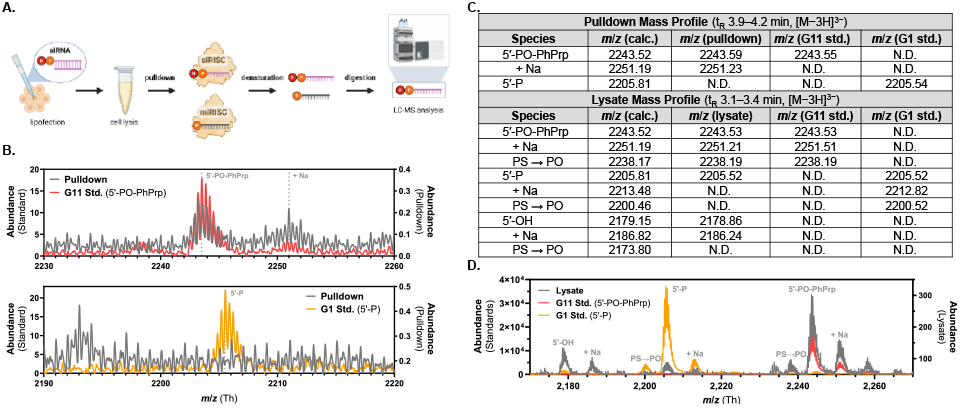
(**A**) Schematic of RISC pulldown method. (**B**) Aligned mass spectra of **G11** pulldown fraction, **G11** standard, and **G1** standard. (**C**) Detected masses and corresponding species in mass analyses. (**D**) Aligned mass spectra of **G11** lysate, **G11** standard, and **G1** standard

Because the bead-bound RISC fraction predominantly comprises endogenous microRNA-loaded RISC,^37^ this background can interfere with subsequent analysis. To selectively isolate RISC-loaded siRNA guide strands (or their metabolites), the pulldown fraction was treated with aqueous piperidine (10% v/v) at 95 °C for 90 min, decanted from the beads, lyophilized to remove piperidine, and precipitated with cold isopropanol.^38^ This procedure digests endogenous RNAs and removes some protein without degrading 5′-POR siRNA guide strands. Following sample preparation, the **G11** pulldown fraction was analyzed by LC-MS (see supporting information) and compared to 5′-PO-PhPrp guide strand (**G11**) and 5′-P (**G1**) standards. The pulldown mass spectrum (**Fig. 5B**) revealed a signal profile (isotopomer envelope) that precisely matches the **G11** standard in its most abundant [M−3H]^3−^ protonation state (**Fig. 5C**), indicating that the **G11** loads intact into RISC in cellula and persists for at least 48 hours. A less abundant signal corresponding to the predicted **G11** sodium adduct was also observed. Importantly, no signal corresponding to the **G1** standard was observed above the limit of detection, suggesting that an intact 5′-PO-PhPrp guide strand, rather than its potential 5′-P metabolite, is the principal active species in RISC.

To explore the total intracellular fraction of **G11** (comprising mostly unloaded siRNA),^39,40^ the **G11** lysate was analyzed via LC-MS and compared to **G11** and **G1** standards (see supporting information). For sample preparation, lysate was denatured at 95 °C for 20 min, extracted with acidchloroform-phenol, depleted of large nucleic acids with solid-phase reversible immobilization beads, and precipitated with cold isopropanol; **G11** and **G1** standards were likewise prepared after spiking into naive lysate. The signal profile of the **G11** lysate (**Fig. 5D**) principally corresponded to the **G11** standard, including minor sodium adduct and PS to PO degradant signals (**Fig. 5C**). However, some 5′-OH and trace 5′-P signals were also observed, suggesting minor intracellular metabolism of **G11** over 48 h. Overall, these results indicate that **G11** is mostly stable and RISC-compatible in cellula.

### Structure determination of 5′-POR guide strands in AGO2

To investigate how AGO2 engages 5′-POR siRNA guide strands, we prepared fully chemically modified oligonucleotides bearing 5′-PO-PhPrp (**ON10**), 5′-PO-PhPr (**ON11**), and 5′-PO-Me (**ON12**), as well as a 5′-P control (**ON13**). Each oligonucleotide was loaded into recombinant AGO2, and the resulting AGO2:oligonucleotide complexes were purified and characterized. Cleavage kinetics of a target RNA by AGO2 loaded with either **ON10** or **ON12** were indistinguishable from AGO2 loaded with the **ON13** control (**Fig. 6A**). By contrast, cleavage guided by **ON11** proceeded at about half the rate (**Fig. 6B**).

**Figure 6.**
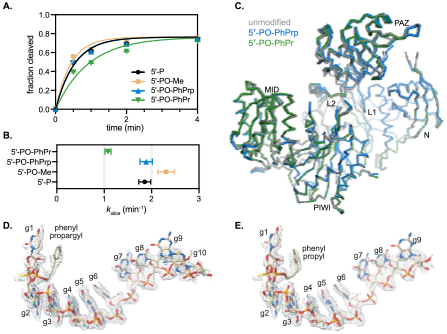
(**A**) Single-turnover target RNA cleavage kinetics by AGO2 loaded with either 5′-P (**ON13**), 5′-PO-Me (**ON12**), 5′-PO-PhPrp (**ON10**), or 5′-PO-PhPr (**ON11**). (**B**) First-order rate constants calculated from data in panel A. Error bars indicate SEM from *n* = 3 trials. (**C**) Superposition of C_a_ backbones of AGO2 loaded with 5′-PO-PhPrp guide (**ON10**) or 5′-PO-PhPr guide (**ON11**) on the equivalent unmodified guide;^41^ AGO2 domains are indicated. (**D**) Experimental F_o_–F_c_ map corresponding to bound **ON11**. (**E**) F_o_–F_c_ map corresponding to bound **ON10**.

We next crystallized the AGO2:**ON10** and AGO2:**ON11** complexes. Crystallization and diffraction properties were similar to AGO2 with an unmodified guide strand (**Table S10**).^42^ Diffraction analysis showed that the backbone conformation of AGO2 was nearly identical to that observed with a fully unmodified guide strand of the same sequence (**Fig. 6C**),^41^ consistent with the notion that crystallization captures a largely invariant AGO2 conformation.^41,43^ The guide strands themselves were also similarly placed as in previous AGO2 structures, with a notable exception of guide (g) nucleotides g8 to g10, which are typically disordered in unmodified structures,^41,42^ but showed well-ordered electron density in the modified RNA structures (**Fig. 6D, E**). The improved order may be due to a 2′-*O*-Me modification on g8, which appears to form favorable hydrophobic interactions with M364 of AGO2.

### A T-shaped π-stacking network stabilizes 5′ phenyl groups

The 5′-PO-PhPrp and 5′-PO-PhPr modifications were resolved clearly in the experimental electron-density maps (**Fig. 6D, E**). Both 5′-P mimics occupy a similar position, interacting with a hydrophobic patch of the AGO2 surface that is directly above the 5′-P binding pocket (**Fig. 7A**). The phenyl rings lie perpendicular to the side chain of Y529 at a distance consistent with T-shaped π-stacking interactions. Through this contact, the phenyl groups join a π-stacking network that includes Y529, F811, and F815 (**Fig. 7B, C**). The presence of these 5′ modifications coincides with a ∼1 Å displacement of the g1 nucleobase relative to the unmodified guide structure. This shift enables stacking between the phenyl groups and Y529 while diminishing stacking between Y529 and the g1 base, which forms a staggered π–π stack in unmodified AGO2:RNA complexes.^13^ Thus, the phenyl substituents partially substitute for interactions normally contributed by the g1 nucleobase.

**Figure 7.**
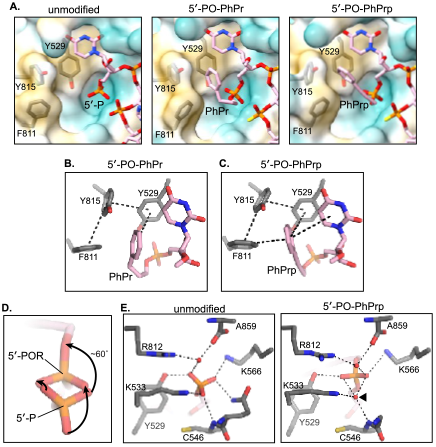
(**A**) Close-up view of the 5′-P binding pocket for unmodified, 5′-PO-PhPr, and 5′-PO-PhPrp structures. The protein surface is colored by hydrophobicity, where cyan indicates hydrophilic and tan indicates hydrophobic. (**B**) The T-shaped π-stacking network surrounding the PhPr group. Dashed lines indicate T-shaped π-stacking interactions. (**C**) The T-shaped π-stacking network surrounding the PhPrp group. (**D**) Superposition of unmodified 5′-P and 5′-PO-PhPrp groups. Viewpoint shows the three non-bonding oxygen atoms in the unmodified, with the P—O^5′^ bond behind, extending away from the viewer. Arrows indicate the direction of rearrangement from unmodified to modified. (**E**) Hydrogen bonding network for the unmodified 5′-P (left) and 5′-POR (right) phosphate groups. Dashed lines indicate hydrogen bonds. Red spheres indicate water molecules. Arrowhead indicates position of the water molecule unique to the 5′-POR structures.

The phenylpropargyl substituent engages in additional T-shaped stacking interactions with both F811 and the g1 nucleobase (**Fig. 7C**). These contacts are not predicted for 5′-PO-PhPr, as flexibility in the propyl linker allows the phenyl group to refine to a position rotated by ∼15° relative to the 5′-PO-PhPrp. The rigid alkyne linker in the phenylpropargyl substituent, therefore, uniquely orients the phenyl ring to establish a four-way stacking network involving Y529, F811, and the g1 nucleobase. These enhanced interactions may underlie the improved silencing potency of the 5′-PO-PhPrp modification compared to the 5′-PO-PhP modification (**Fig. 3**).

### An ordered water molecule compensates for displacement of the 5′-POR

In unmodified AGO2:RNA, the 5′-P forms an extensive hydrogen-binding network involving all three nonbridging oxygen atoms.^13^ These oxygens interact with the main chain amide of C546, side chains of Q545, Y529, K533, K570, and K566, and an ordered water molecule. Remarkably, despite the addition of the organic substituents, many of these interactions are preserved in the 5′-POR structures. Accommodation of the substituent requires the modified 5′-P to shift by ∼1 Å relative to the unmodified position and rotate by 60° around the dihedral angle *β*. This movement effectively inverts the phosphate orientation such that two non-bridging oxygen atoms remain near their unmodified positions, while the third oxygen atom, featuring the substituent, flips to the opposite side of the phosphorus (**Fig. 7D**). The resulting displacement creates a gap between the 5′-POR and the main chain amide of C546. This void is occupied by an ordered water molecule that bridges the C546 amide and K533 to a non-bridging oxygen in the 5′-POR, forming a new hydrogen-bonding network with nearly the same connectivity as in the unmodified structure (**Fig. 7E**). Thus, reorganization of the 5′-P hydrogen-bonding network through the ordered water molecule appears to compensate for interactions lost upon displacement of the 5′-P.

Overall, these structures reveal a distinct guide strand 5′-end binding mode in which the 5′-P mimic substituents remodel local contacts while preserving overall binding stability, offering mechanistic insight into how 5′-P mimics that enhance siRNA stability can still provide functional efficacy.

## CONCLUSIONS

In this study, we developed facile “on-support” methods to access modified siRNA guide strands with three previously uncharacterized classes of 5′-P mimics (5′-POR, 5′-PSR, and 5′-MsPA). These methods are applicable to synthetic oligonucleotides generally, affording exploration of their properties in new contexts. We prepared a diverse panel of thirtyfive 5′-POR guide strands and identified two leads (5′-PO-Me and 5′-PO-PhPrp) with high RISC compatibility. Likewise, we demonstrated high RISC compatibility for 5′-PS, 5′-PS-PhPrp, and 5′-MsPA. Collectively, these findings show that AGO2 accommodates diverse chemical modifications of 5′-P, including substituents on a non-bridging oxygen or sulfur, or replacement of a non-bridging oxygen with sulfur or the mesylimino group. The crystal structure of a 5′-PO-PhPrp guide strand bound to AGO2 revealed new π–π interactions between the rigid phenylpropargyl substituent and an aromatic network adjacent to the 5′ nucleotide binding site. In parallel, rearrangement of the hydrogen-binding network within the 5′-P binding pocket allows AGO2 to ac-commodate organic substituents attached to the 5′-P of guide strands. These structural insights into the SAR of RISC compatibility provide a foundation for iterative optimization of 5′-POR anchoring within RISC using rationally designed new substituents. We subsequently used our novel 5′-P mimics to identify structural determinants of in vitro oligonucleotide susceptibility to two known degradative enzymes, alkaline phosphatase and XRN1 (a major 5′-exonuclease). Although the phosphatase is highly selective for 5′-P guide strands, XRN1 is more sensitive to the net charge of the 5′-P mimic than to its steric demand, selectively degrading dianionic 5′-P mimics of various sizes and shapes. The results of our phosphatase susceptibility assays demonstrate that a carbon–phosphorus bond is not an essential feature of phosphatase-resistant 5′-P mimics. Additionally, the trends identified in both enzyme assays may be used to predict the enzymatic susceptibility of potential new 5′-P mimics and extend metabolic stabilization to other classes of synthetic oligonucleotides (provided that such 5′-P mimics do not disrupt any functional supramolecular interactions). However, the complex composition of endolysosomal and cytosolic environments does not permit generalization of these in vitro observations to the in vivo metabolic stability of siRNA. Finally, we demonstrated that intact 5′-PO-PhPrp siRNA persists for at least 48 h and loads into RISC in cellula. Because the lead 5′-POR and 5′-PSR variants stabilize guide strands against both alkaline phosphatase and XRN1 in vitro while preserving potent RNAi activity in cells, future exploration of their properties in vivo are also warranted.

## ASSOCIATED CONTENT

### Data Availability Statement

Coordinate data for the crystal structures of the AGO2:**ON10** and AGO2:**ON11** complexes have been deposited in the Protein Data Bank (https://www.rcsb.org) with the identifiers 9OBD and 9OBE, respectively.

### Supporting Information

The following file is available on https://www.biorxiv.org.

- Experimental procedures, authentication of oligonucleotides, supplemental silencing activity data, diffraction data collection information and refinement statistics (PDF)

### Author Contributions

K.Y. and A.K. conceived the project. T.C.B. and K.Y. designed 5′-POR. T.C.B. synthesized POR and conducted enzyme stability assays. T.C.B., S.H., and E.L. designed and conducted AGO2 pulldown experiments. L.F.R.G. and I.J.M. determined crystal structures and conducted RISC kinetics analysis. T.C.B., N.Y., J.C., and E.L. conducted in vitro potency assays. N.M. and D.E. supported solid-phase synthesis methods and mass analyses. J.K.W. and A.W. conceived and supported the 5′-MsPA modification scheme. T.C.B., K.Y., and A.K. wrote the manuscript. All authors have given approval to the final version of the manuscript.

### Funding Sources

NIH R35 GM131839

NIH S10 OD20012

NIH S10 OD036329

University of Oxford subaward of United Kingdom Medical Research Council

### Notes

A.K. is a co-founder, scientific advisory board member, and shareholder of Atalanta Therapeutics, as well as a founder of Comanche Pharmaceuticals and on the scientific advisory board of Advirna, Alys Pharmaceuticals, and Prime Medicine.

## ACKNOWLEDGMENT

We thank Dr. A. Ali for maintaining NMR instruments and Dr. S. Nguyen for HRMS analysis. The authors would also like to thank Dr. E. Haberlin for copy editing. Some portions of figures were created in BioRender.com.

