## Supporting Information for "Organyl 5′‑Phosphates in siRNA Guide Strands: Structure–Function Relationships Governing Anchoring in Argonaute 2 and Metabolic Stability"

### Table of Contents


| Section | Page |
| --- | --- |
| Oligonucleotide Synthesis and Modification | S2 |
| Oligonucleotide Characterization and Purification | S6 |
| Cell Culture | S7 |
| In Cellula Silencing Efficacy and Potency Assays | S7 |
| In Vitro Enzyme Susceptibility Assays | S13 |
| In Cellula Metabolism and RISC Loading Studies | S16 |
| Biochemical and Structural Analyses | S18 |
| Oligonucleotide Authentication | S20 |
| References | S22 |

### Oligonucleotide Synthesis and Modification

***General synthesis conditions***. 3′-Cholesterol (3′-Chol) oligonucleotides, which feature a cholesterol moiety bound to the 3′-end via a tetraethylene glycol linker, were synthesized on Cholesterol 3′-lcaa solid support (ChemGenes); all other oligonucleotides were synthesized on UnyLinker 500 Å solid support (ChemGenes). All oligonucleotides were synthesized at a 5–10 µmol scale using a MerMade 12 synthesizer (BioAutomation). Stepwise 3′-to-5′ synthesis was conducted via the phosphoramidite synthesis cycle, comprising (1) deblocking, (2) coupling, (3) capping, and (4) oxidation or sulfurization. Deblocking (detritylation) was performed with deblocking reagent (Glen Research) comprising 3% w/v trichloroacetic acid in dry $CH_2Cl_2$. For the coupling step, standard phosphoramidite reagents (ChemGenes) consisting of 5′-*O*-dimethoxytrityl nucleoside 3′-(*N*,*N*-diisopropyl-*O*-cyanoethyl) phosphoramidite with a protected nucleobase ($A^{Bz}$, $C^{Ac}$, $G^{Ac}$, or U) and modified ribose (2′-*O*-Me or 2′-F) were used in dry MeCN (0.1 M); all coupling times were 10 min. For 5′-(*E*)-vinylphosphonate (5′-*E*-VP) oligonucleotides, 5′-bis(pivaloyloxymethyl)-*E*-vinylphosphonate-2′-*O*-methyl-uridine 3′-(*N*,*N*-diisopropyl-*O*-cyanoethyl) phosphoramidite (Hongene Biotechnology) was used for the terminal coupling. Coupling activation was performed with activator reagent (Glen Research) comprising 0.25 M 5-ethylthio-1*H*-tetrazole (ETT) in dry MeCN. Capping (5′-*O*-acetylation) of unreacted shortmers was achieved with capping reagent A comprising 1-methylimidazole/2,6-lutidine/MeCN (2:3:5 v/v/v) and capping reagent B comprising 20% v/v $Ac_2O$ in MeCN (Glen Research). Oxidation or sulfurization of the intermediate phosphite was performed with oxidation reagent (ChemGenes) comprising 0.05 M iodine in pyridine/$H_2O$ (9:1 v/v), or sulfurization reagent (ChemGenes) comprising 0.1 M ((dimethylaminomethylidene)amino)-3*H*-1,2,4-dithiazoline-3-thione (DDTT) in dry pyridine/MeCN (9:1 v/v), for 2.5 min. All oligonucleotides with a dimethoxytrityl group on the terminal hydroxyl (5′-OH) after synthesis were deblocked ("trityl-off"). Synthesis was monitored via inline detection of the dimethoxytritylium byproduct during each deblocking step.

After any 5′-end modification (see below), oligonucleotides were cleaved from solid support and deprotected via the ammonia-methylamine method (conc. $NH_4OH$ and 40% w/v $MeNH_2$ in $H_2O$, 1:1 v/v; rt, 2 h). 5′-*E*-VP oligonucleotides were deprotected in 3% v/v $Et_2NH$ in conc. $NH_4OH$ for 20 h at 35 °C. Subsequently, solid support was filtered off, and volatile bases were removed by lyophilization to yield the crude oligonucleotide product.

***General modification conditions.*** Unless otherwise stated, all modifications of on-support 5′-OH oligonucleotides (see *General synthesis conditions*) were performed in anhydrous conditions under argon at room temperature; reagents were sourced from Sigma-Aldrich, Acros, or TCI at reagent-grade purity. Dried ACS-grade solvents were taken from a PureSolv MD7 solvent purification system (Innovative Technology) under argon and stored over drying traps (ChemGenes) until use. Liquid reagents were dried overnight on 3Å molecular sieves; aliquots of solid reagents were taken directly. Standard activator reagent, oxidation reagent, and sulfurization reagent were used (see *General synthesis conditions*). Dryness of materials (<20 ppm $H_2O$) were confirmed via Karl Fisher titration using a Metrohm 899 Coulometer. Reactions were performed in fritted peptide synthesis vessels (Kemtech) capped with airtight rubber septa. Synthesis vessels and on-support 5′-OH oligonucleotides were washed thrice with dry MeCN under argon prior to use.

**Scheme S1**. 5′-P and 5′-PS modification.

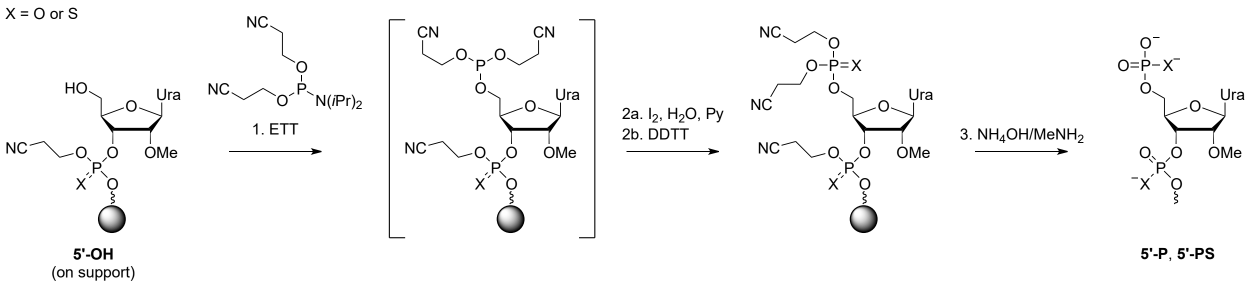


***5′-P and 5′-PS modification.*** For step 1 (**Scheme S1**), on-support 5′-OH oligonucleotide was coupled for 10 min with 0.1 M *O*,*O*′-bis(cyanoethyl)-*N*,*N*-diisopropyl phosphoramidite (ChemGenes) in MeCN on the MerMade 12 synthesizer, followed by standard oxidation (step 2a) for 5′-P modification or standard sulfurization (step 2b) for 5′-PS modification (see *General synthesis conditions*).

**Scheme S2**. 5′-POR and 5′-PSR modification.

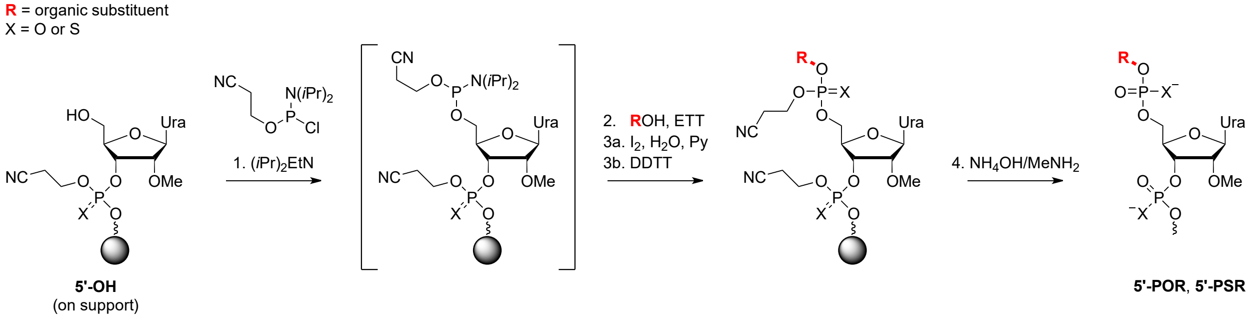


**Figure S1.** Index of 5′-POR and 5′-PSR substituents (R).

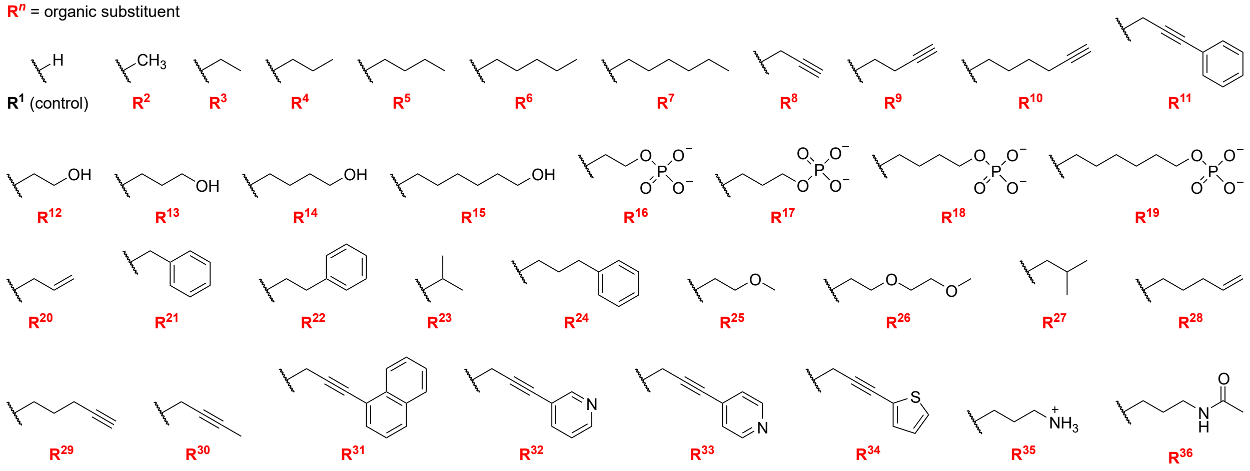


These substituent indices correspond to the guide strand IDs in the 5′-POR screening panels (**Table S1, S3**).

***General 5′-POR and 5′-PSR modification.*** For step 1 (**Scheme S2**), MeCN (1.2 mL) was added via syringe to a synthesis vessel containing on-support 5′-OH oligonucleotide (2.5 µmol, 1.0 eq), followed by *i*$Pr_2$EtN (174 µL, 1,000 µmol, 400 eq). Neat *N*,*N*-diisopropyl-*O*-cyanoethyl chlorophosphoramidite (ChemGenes; 112 µL, 500 µmol, 200 eq) was then added dropwise via syringe. The reaction was gently agitated for 1 h, after which the supernatant was purged, and the solid support was washed with MeCN (~3 mL). For step 2, an aliquot of neat alcohol (1,000 µmol, 400 eq) with the desired substituent (**$R^3$–$R^{11}$**, **$R^{20}$–$R^{34}$**; Fig. S1) was added via syringe over the solid support, followed by activator reagent (2.5 mL). The reaction was agitated gently for 2 h, then supernatant was purged, and solid support was washed with MeCN (~3 mL). For oxidation (step 3a) or sulfurization (step 3b), excess oxidation or sulfurization reagent (~2.5 mL) was added via pipette under room air. After 1 min of oxidation or 2.5 min of sulfurization, supernatant was purged and solid support was washed with MeOH, yielding on-support 5′-POR or 5′-PSR oligonucleotides. To broaden the functional group tolerance of **Scheme S2** or reduce synthetic steps, the synthetic approach was modified for **$R^2$**, **$R^{12}$–$R^{19}$**, and **$R^{35}$–$R^{36}$** (see below).

**Figure S2.** "Spacer" phosphoramidite reagents for 5′-POR modification.

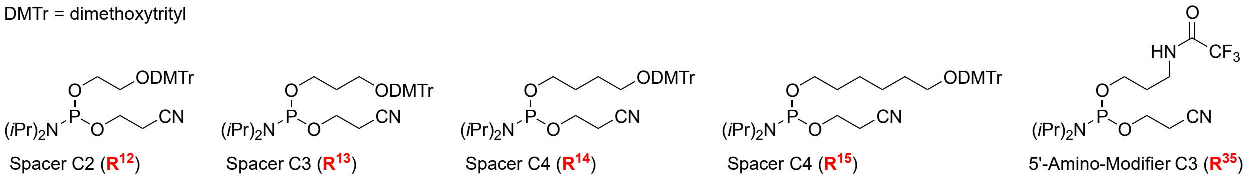


For **$R^{12}$–$R^{15}$** and **$R^{35}$** (Fig. S1), the on-support 5′-OH oligonucleotide was coupled to commercial "spacer" and "5′-amino modifier" phosphoramidites (Glen Research; **Fig. S2**) and oxidized on the MerMade 12 synthesizer (see *General oligonucleotide synthesis*), yielding on-support 5′-POR oligonucleotides. These phosphoramidites protect the substituents with acid-labile dimethoxytrityl (DMTr) or trifluoroacetyl groups, which are readily deblocked after synthesis. This approach is more efficient than using protected diols and amino-alcohols in **Scheme S2**.

**Scheme S3**. 5′-PO-Me modification.

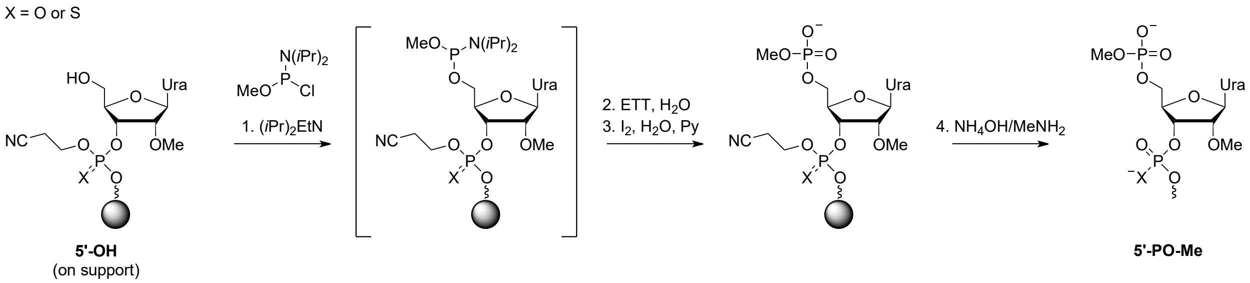


***5′-PO-Me modification.*** For **$R^2$** (Me), an efficient approach featuring commercial *N*,*N*-diisopropyl-*O*-methyl chlorophosphoramidite reagent was used (**Scheme S3**). In step 1, MeCN (1.2 mL) was added via syringe to the synthesis vessel containing on-support 5′-OH oligonucleotide (2.5 µmol, 1.0 eq), followed by *i*$Pr_2$EtN (174 µL, 1,000 µmol, 400 eq). The synthesis vessel was swirled to uniformly disperse the *i*$Pr_2$EtN in solution, then neat *N*,*N*-

diisopropyl-*O*-methyl chlorophosphoramidite (ChemGenes; 97 µL, 500 µmol, 200 eq) was added dropwise via syringe. The reaction was agitated gently for 3 h, after which the supernatant was purged and the solid support was washed with MeCN (~3 mL). For step 2, activator reagent (2.5 mL) was added via syringe, followed by excess ACS-grade $H_2O$ (300 µL). The reaction was agitated gently for 15 min, then supernatant was purged and solid support was washed with MeCN (~3 mL). Oxidation (step 3a) or sulfurization (step 3b) was performed as above (see *General 5′-POR and 5′-PSR modification*), yielding the on-support 5′-PO-Me oligonucleotide. For reasons not evident to the authors, these conditions proved unsuitable for 5′-PS-Me modification.

**Scheme S4**. Phosphorylation of select 5′-POR modifications.

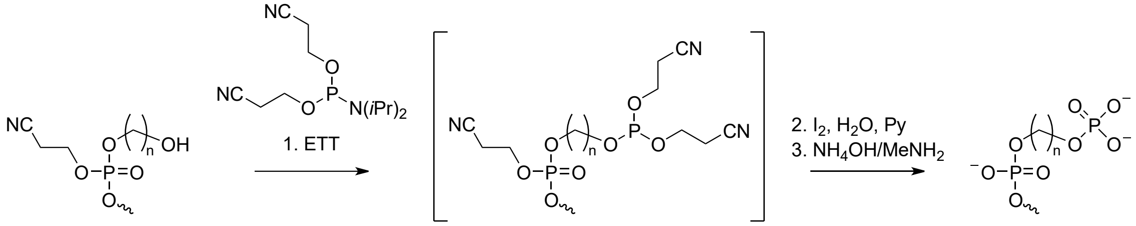


**5'-POR$^{12–15}$** (on support) **5'-POR$^{16–19}$**

**Scheme S5**. *N*-Acetylation of select 5′-POR modification.

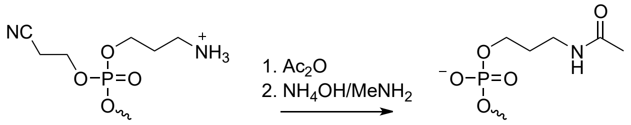


**5'-POR$^{35}$** (on support) **5'-POR$^{36}$**

***Secondary 5′-POR derivatization.*** Phosphate (**R$^{16}$**–**R$^{19}$**) and *N*-acetyl (**R$^{36}$**) modifications were derivatized from ω-hydroxy (**R$^{12}$**–**R$^{15}$**) and ω-amine (**R$^{35}$**) precursors, respectively (see *General 5′-POR and 5′-PSR modification*). The on-support precursor modifications were deblocked on the MerMade 12 synthesizer. Subsequently, **R$^{12}$**–**R$^{15}$** were phosphorylated with *O*,*O*′-bis(cyanoethyl)-*N*,*N*-diisopropyl phosphoramidite and oxidized (see *5′-P and 5′-PS modification*), yielding on-support **R$^{16}$**–**R$^{19}$** products, respectively (**Scheme S4**). **R$^{35}$** was acetylated (capped) via standard methods (see *General synthesis conditions*), yielding the on-support **R$^{36}$** product (**Scheme S5**). Derivatization of 5′-PSR modifications was not attempted within the scope of this work.

**Scheme S6**. 5′-MsPA modification.

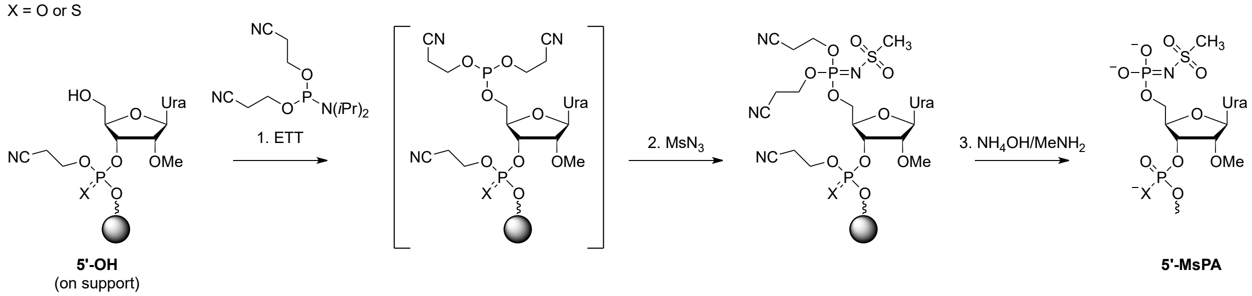


***5′-MsPA modification.*** For step 1 (**Scheme S6**), on-support 5′-OH oligonucleotide (5.0 µmol, 1 eq) was coupled for 10 min with 0.1 M *O*,*O′*-bis(cyanoethyl)-*N*,*N*-diisopropyl phosphoramidite (ChemGenes) in MeCN on the MerMade 12 synthesizer (see *General synthesis conditions*). Subsequently, the on-support phosphoramidite intermediate was oxidized for 18 min (3 min wait, 5 × 3 min vacuum pulse) with freshly prepared 0.5 M mesyl azide in MeCN (700 µL, 350 µmol, 70 eq) on the MerMade 12 synthesizer, yielding on-support 5′-MsPA modification.

The mesyl azide solution was prepared as follows. Mesyl chloride (193 µL, 2.5 mmol, 1.0 eq) was added to dry MeCN (5.0 mL) in a conical tube over ice, followed by $NaN_3$ (195 mg, 3.0 mmol, 1.2 eq); some $NaN_3$ remained undissolved. The reaction was then gently agitated overnight at room temperature, then the supernatant was decanted and used directly.

**Oligonucleotide Characterization and Purification**

Identity of all oligonucleotides was determined by LC-MS analysis on an Agilent 6530 Accurate Mass Q-TOF LC-MS instrument using the following conditions: AdvanceBio Oligonucleotide C18 LC column (Agilent), 60 °C, 0.85 mL/min; buffer A: 50 mM hexafluoroisopropanol, 10 mM $Et_3N$ in $H_2O$; buffer B: 50 mM hexafluoroisopropanol, 10 mM $Et_3N$ in MeOH; gradient profile: 5% B for 1.6 min, 5–100% B for 5.5 min, 100% B for 1.6 min, 5% B for 1.6 min; $A_{260}$ peak monitoring. ESI-MS parameters were as follows: negative-ion centroid mode, 4,000 V capillary potential, 100–3,200 m/z range, 2 Hz scan rate. Mass spectra were extracted and deconvoluted using Agilent MassHunter software. The calculated and observed masses of all oligonucleotides used in this work are listed in **Table S11**.

All crude oligonucleotides were purified via HPLC on an Agilent 1260 Infinity instrument using the following conditions: Hamilton PRP-C18 column, 60 °C, 30 mL/min; buffer A: 50 mM NaOAc, 5% v/v MeCN in $H_2O$; buffer B: 100% MeCN; gradient profile: 0% B for 5 min, 0–50% B for 20 min, 100% B for 5 min, 0% B for 5 min (3′-Chol oligonucleotides); buffer A: 100 mM hexafluoroisopropanol, 15 mM $Et_3N$ in $H_2O$; buffer B: 100 mM hexafluoroisopropanol, 15 mM $Et_3N$ in MeOH; gradient profile: 18% B for 5 min, 18%–35% B for 30 min, 100% B for 5 min, 18% B for 5 min (all other oligonucleotides); $A_{260}$, $A_{230}$ peak monitoring. Fractions were analyzed via LC-MS (see above) and pooled. Sodium salt exchange was conducted by anion-exchange HPLC on an Agilent 1260 Infinity instrument using the following conditions: Cytiva

Source 15Q column, 50 °C, 45 mL/min; buffer A: 10 mM NaOAc, 10% v/v MeCN in $H_2O$; buffer B: 1.0 M NaBr, 10 mM NaOAc, 10% v/v MeCN in $H_2O$; gradient profile: 30% B for 3 min, 30–80% B for 15 min, 100% B for 5 min, 30% B for 5 min. Desalting was conducted by size-exclusion chromatography on an AKTA P-920 FPLC instrument (Amersham) using the following conditions: Cytiva Sephadex G25 column (120 mL), 20 mL/min $H_2O$, $A_{260}$ and conductivity peak monitoring. Desalted oligonucleotide stocks were quantified by $A_{260}$ on a NanoDrop 1000 spectrophotometer (Thermo Scientific).

To prepare siRNA duplexes, equimolar aliquots of a guide strand and its complementary passenger strand were combined in nuclease-free $H_2O$, denatured at 95 °C for 5 min, then annealed by gradual cooling to room temperature (~3 h). Duplex formation was confirmed via non-denaturing PAGE on Novex 20% TBE gels (Invitrogen) followed by staining with SYBR Gold (ThermoFisher) and visualization on a Bio-Rad Gel Doc Go fluorescence gel imager.

### Cell Culture

HeLa cells (ATCC) from cryo-storage were seeded into culture plates in Dulbecco's modified Eagle medium (DMEM; Gibco) supplemented with 10% fetal bovine serum (FBS; Gibco). Cells were incubated at 37 °C and 5% $CO_2$ and passaged after growth to 80% confluency; unused cells were discarded by the tenth passage. For all seeding aliquots, cell density and percent viability were quantified in technical duplicate on a Cellometer Auto 1000 cell counter (Nexcelcom) with 0.4% trypan blue staining (Gibco).

### In Cellula Silencing Efficacy and Potency Assays

**Table S1**. Primary panel of cholesterol-conjugated siRNAs for silencing efficacy screen.

| Duplex ID | Target | Strand ID | Sequence (5′ to 3′) |
|---|---|---|---|
| *Positive Control* | | | |
| **D1** | *HTT* | **G1** | P(mU)#(fU)#(mA)(fA)(mU)(fC)(mU)(fC)(mU)(fU)(mU)(fA)(mC)#(fU)#(mG)#(fA)#(mU)#(fA)#(mU)#(fA) |
| | | **P1** | (fC)#(mA)#(fG)(mU)(fA)(mA)(fA)(mG)(fA)(mG)(fA)(mU)(fU)#(mA)#(fA)Chol |
| *Test Articles (Primary Panel)* | | | |
| **D2** | *HTT* | **G2** | methyl-P(mU)#(fU)#(mA)(fA)(mU)(fC)(mU)(fC)(mU)(fU)(mU)(fA)(mC)#(fU)#(mG)#(fA)#(mU)#(fA)#(mU)#(fA) |
| | | **P1** | (fC)#(mA)#(fG)(mU)(fA)(mA)(fA)(mG)(fA)(mG)(fA)(mU)(fU)#(mA)#(fA)Chol |
| **D3** | *HTT* | **G3** | ethyl-P(mU)#(fU)#(mA)(fA)(mU)(fC)(mU)(fC)(mU)(fU)(mU)(fA)(mC)#(fU)#(mG)#(fA)#(mU)#(fA)#(mU)#(fA) |
| | | **P1** | (fC)#(mA)#(fG)(mU)(fA)(mA)(fA)(mG)(fA)(mG)(fA)(mU)(fU)#(mA)#(fA)Chol |
| **D4** | *HTT* | **G4** | propyl-P(mU)#(fU)#(mA)(fA)(mU)(fC)(mU)(fC)(mU)(fU)(mU)(fA)(mC)#(fU)#(mG)#(fA)#(mU)#(fA)#(mU)#(fA) |
| | | **P1** | (fC)#(mA)#(fG)(mU)(fA)(mA)(fA)(mG)(fA)(mG)(fA)(mU)(fU)#(mA)#(fA)Chol |
| **D5** | *HTT* | **G5** | butyl-P(mU)#(fU)#(mA)(fA)(mU)(fC)(mU)(fC)(mU)(fU)(mU)(fA)(mC)#(fU)#(mG)#(fA)#(mU)#(fA)#(mU)#(fA) |
| | | **P1** | (fC)#(mA)#(fG)(mU)(fA)(mA)(fA)(mG)(fA)(mG)(fA)(mU)(fU)#(mA)#(fA)Chol |
| **D6** | *HTT* | **G6** | pentyl-P(mU)#(fU)#(mA)(fA)(mU)(fC)(mU)(fC)(mU)(fU)(mU)(fA)(mC)#(fU)#(mG)#(fA)#(mU)#(fA)#(mU)#(fA) |
| | | **P1** | (fC)#(mA)#(fG)(mU)(fA)(mA)(fA)(mG)(fA)(mG)(fA)(mU)(fU)#(mA)#(fA)Chol |
| **D7** | *HTT* | **G7** | hexyl-P(mU)#(fU)#(mA)(fA)(mU)(fC)(mU)(fC)(mU)(fU)(mU)(fA)(mC)#(fU)#(mG)#(fA)#(mU)#(fA)#(mU)#(fA) |

| | | | |
|---|---|---|---|
| | | P1 | (fC)#(mA)#(fG)(mU)(fA)(mA)(fA)(mG)(fA)(mG)(fA)(mU)(fU)#(mA)#(fA)Chol |
| D8 | *HTT* | G8 | propargyl-P(mU)#(fU)#(mA)(fA)(mU)(fC)(mU)(fC)(mU)(fU)(mU)(fA)(mC)#(fU)#(mG)#(fA)#(mU)#(fA)#(mU)#(fA) |
| | | P1 | (fC)#(mA)#(fG)(mU)(fA)(mA)(fA)(mG)(fA)(mG)(fA)(mU)(fU)#(mA)#(fA)Chol |
| D9 | *HTT* | G9 | 3-butynyl-P(mU)#(fU)#(mA)(fA)(mU)(fC)(mU)(fC)(mU)(fU)(mU)(fA)(mC)#(fU)#(mG)#(fA)#(mU)#(fA)#(mU)#(fA) |
| | | P1 | (fC)#(mA)#(fG)(mU)(fA)(mA)(fA)(mG)(fA)(mG)(fA)(mU)(fU)#(mA)#(fA)Chol |
| D10 | *HTT* | G10 | 5-hexynyl-P(mU)#(fU)#(mA)(fA)(mU)(fC)(mU)(fC)(mU)(fU)(mU)(fA)(mC)#(fU)#(mG)#(fA)#(mU)#(fA)#(mU)#(fA) |
| | | P1 | (fC)#(mA)#(fG)(mU)(fA)(mA)(fA)(mG)(fA)(mG)(fA)(mU)(fU)#(mA)#(fA)Chol |
| D11 | *HTT* | G11 | phenylpropargyl-P(mU)#(fU)#(mA)(fA)(mU)(fC)(mU)(fC)(mU)(fU)(mU)(fA)(mC)#(fU)#(mG)#(fA)#(mU)#(fA)#(mU)#(fA) |
| | | P1 | (fC)#(mA)#(fG)(mU)(fA)(mA)(fA)(mG)(fA)(mG)(fA)(mU)(fU)#(mA)#(fA)Chol |
| D12 | *HTT* | G12 | 2-hydroxyethyl-P(mU)#(fU)#(mA)(fA)(mU)(fC)(mU)(fC)(mU)(fU)(mU)(fA)(mC)#(fU)#(mG)#(fA)#(mU)#(fA)#(mU)#(fA) |
| | | P1 | (fC)#(mA)#(fG)(mU)(fA)(mA)(fA)(mG)(fA)(mG)(fA)(mU)(fU)#(mA)#(fA)Chol |
| D13 | *HTT* | G13 | 3-hydroxypropyl-P(mU)#(fU)#(mA)(fA)(mU)(fC)(mU)(fC)(mU)(fU)(mU)(fA)(mC)#(fU)#(mG)#(fA)#(mU)#(fA)#(mU)#(fA) |
| | | P1 | (fC)#(mA)#(fG)(mU)(fA)(mA)(fA)(mG)(fA)(mG)(fA)(mU)(fU)#(mA)#(fA)Chol |
| D14 | *HTT* | G14 | 4-hydroxybutyl-P(mU)#(fU)#(mA)(fA)(mU)(fC)(mU)(fC)(mU)(fU)(mU)(fA)(mC)#(fU)#(mG)#(fA)#(mU)#(fA)#(mU)#(fA) |
| | | P1 | (fC)#(mA)#(fG)(mU)(fA)(mA)(fA)(mG)(fA)(mG)(fA)(mU)(fU)#(mA)#(fA)Chol |
| D15 | *HTT* | G15 | 6-hydroxyhexyl-P(mU)#(fU)#(mA)(fA)(mU)(fC)(mU)(fC)(mU)(fU)(mU)(fA)(mC)#(fU)#(mG)#(fA)#(mU)#(fA)#(mU)#(fA) |
| | | P1 | (fC)#(mA)#(fG)(mU)(fA)(mA)(fA)(mG)(fA)(mG)(fA)(mU)(fU)#(mA)#(fA)Chol |
| D16 | *HTT* | G16 | 2-phosphonooxyethyl-P(mU)#(fU)#(mA)(fA)(mU)(fC)(mU)(fC)(mU)(fU)(mU)(fA)(mC)#(fU)#(mG)#(fA)#(mU)#(fA)#(mU)#(fA) |
| | | P1 | (fC)#(mA)#(fG)(mU)(fA)(mA)(fA)(mG)(fA)(mG)(fA)(mU)(fU)#(mA)#(fA)Chol |
| D17 | *HTT* | G17 | 3-phosphonooxypropyl-P(mU)#(fU)#(mA)(fA)(mU)(fC)(mU)(fC)(mU)(fU)(mU)(fA)(mC)#(fU)#(mG)#(fA)#(mU)#(fA)#(mU)#(fA) |
| | | P1 | (fC)#(mA)#(fG)(mU)(fA)(mA)(fA)(mG)(fA)(mG)(fA)(mU)(fU)#(mA)#(fA)Chol |
| D18 | *HTT* | G18 | 4-phosphonooxybutyl-P(mU)#(fU)#(mA)(fA)(mU)(fC)(mU)(fC)(mU)(fU)(mU)(fA)(mC)#(fU)#(mG)#(fA)#(mU)#(fA)#(mU)#(fA) |
| | | P1 | (fC)#(mA)#(fG)(mU)(fA)(mA)(fA)(mG)(fA)(mG)(fA)(mU)(fU)#(mA)#(fA)Chol |
| D19 | *HTT* | G19 | 6-phosphonooxyhexyl-P(mU)#(fU)#(mA)(fA)(mU)(fC)(mU)(fC)(mU)(fU)(mU)(fA)(mC)#(fU)#(mG)#(fA)#(mU)#(fA)#(mU)#(fA) |
| | | P1 | (fC)#(mA)#(fG)(mU)(fA)(mA)(fA)(mG)(fA)(mG)(fA)(mU)(fU)#(mA)#(fA)Chol |
| D20 | *HTT* | G20 | allyl-P(mU)#(fU)#(mA)(fA)(mU)(fC)(mU)(fC)(mU)(fU)(mU)(fA)(mC)#(fU)#(mG)#(fA)#(mU)#(fA)#(mU)#(fA) |
| | | P1 | (fC)#(mA)#(fG)(mU)(fA)(mA)(fA)(mG)(fA)(mG)(fA)(mU)(fU)#(mA)#(fA)Chol |
| D21 | *HTT* | G21 | benzyl-P(mU)#(fU)#(mA)(fA)(mU)(fC)(mU)(fC)(mU)(fU)(mU)(fA)(mC)#(fU)#(mG)#(fA)#(mU)#(fA)#(mU)#(fA) |
| | | P1 | (fC)#(mA)#(fG)(mU)(fA)(mA)(fA)(mG)(fA)(mG)(fA)(mU)(fU)#(mA)#(fA)Chol |
| D22 | *HTT* | G22 | 2-phenylethyl-P(mU)#(fU)#(mA)(fA)(mU)(fC)(mU)(fC)(mU)(fU)(mU)(fA)(mC)#(fU)#(mG)#(fA)#(mU)#(fA)#(mU)#(fA) |
| | | P1 | (fC)#(mA)#(fG)(mU)(fA)(mA)(fA)(mG)(fA)(mG)(fA)(mU)(fU)#(mA)#(fA)Chol |
| D23 | *HTT* | G23 | isopropyl-P(mU)#(fU)#(mA)(fA)(mU)(fC)(mU)(fC)(mU)(fU)(mU)(fA)(mC)#(fU)#(mG)#(fA)#(mU)#(fA)#(mU)#(fA) |
| | | P1 | (fC)#(mA)#(fG)(mU)(fA)(mA)(fA)(mG)(fA)(mG)(fA)(mU)(fU)#(mA)#(fA)Chol |

Strand IDs are designated as follows: "G": guide strand; "P": passenger strand. Chemical modifications are designated as follows: "#": phosphorothioate linkage; "f": 2′-deoxy-2′-fluoro; "m": 2′-*O*-methyl; "P": 5′-phosphate; Chol": 3′-cholesterol conjugate (ChemGenes). Any substituents are named using standard chemical nomenclature.

**Table S2**. Unconjugated siRNAs of primary panel hits for silencing potency assay.

| Duplex ID | Target | Strand ID | Sequence (5′ to 3′) |
|---|---|---|---|

| | | | |
|---|---|---|---|
| *Positive Control* | | | |
| **D24** | *HTT* | **G1** | P(mU)#(fU)#(mA)(fA)(mU)(fC)(mU)(fC)(mU)(fU)(mU)(fA)(mC)#(fU)#(mG)#(fA)#(mU)#(fA)#(mU)#(fA) |
| | | **P2** | (fC)#(mA)#(fG)(mU)(fA)(mA)(fA)(mG)(fA)(mG)(fA)(mU)(fU)#(mA)#(fA) |
| *Primary Panel Hits* | | | |
| **D25** | *HTT* | **G2** | methyl-P(mU)#(fU)#(mA)(fA)(mU)(fC)(mU)(fC)(mU)(fU)(mU)(fA)(mC)#(fU)#(mG)#(fA)#(mU)#(fA)#(mU)#(fA) |
| | | **P2** | (fC)#(mA)#(fG)(mU)(fA)(mA)(fA)(mG)(fA)(mG)(fA)(mU)(fU)#(mA)#(fA) |
| **D26** | *HTT* | **G4** | propyl-(mU)#(fU)#(mA)(fA)(mU)(fC)(mU)(fC)(mU)(fU)(mU)(fA)(mC)#(fU)#(mG)#(fA)#(mU)#(fA)#(mU)#(fA) |
| | | **P2** | (fC)#(mA)#(fG)(mU)(fA)(mA)(fA)(mG)(fA)(mG)(fA)(mU)(fU)#(mA)#(fA) |
| **D27** | *HTT* | **G8** | 3-propargyl-P(mU)#(fU)#(mA)(fA)(mU)(fC)(mU)(fC)(mU)(fU)(mU)(fA)(mC)#(fU)#(mG)#(fA)#(mU)#(fA)#(mU)#(fA) |
| | | **P2** | (fC)#(mA)#(fG)(mU)(fA)(mA)(fA)(mG)(fA)(mG)(fA)(mU)(fU)#(mA)#(fA) |
| **D28** | *HTT* | **G11** | phenylpropargyl-P(mU)#(fU)#(mA)(fA)(mU)(fC)(mU)(fC)(mU)(fU)(mU)(fA)(mC)#(fU)#(mG)#(fA)#(mU)#(fA)#(mU)#(fA) |
| | | **P2** | (fC)#(mA)#(fG)(mU)(fA)(mA)(fA)(mG)(fA)(mG)(fA)(mU)(fU)#(mA)#(fA) |
| *Structural Comparandum (***G11*** Analog)* | | | |
| **D29** | *HTT* | **G24** | 3-phenylpropyl-P(mU)#(fU)#(mA)(fA)(mU)(fC)(mU)(fC)(mU)(fU)(mU)(fA)(mC)#(fU)#(mG)#(fA)#(mU)#(fA)#(mU)#(fA) |
| | | **P2** | (fC)#(mA)#(fG)(mU)(fA)(mA)(fA)(mG)(fA)(mG)(fA)(mU)(fU)#(mA)#(fA) |

Strand IDs are designated as follows: “G”: guide strand; “P”: passenger strand. Chemical modifications are designated as follows: “#”: phosphorothioate linkage; “f”: 2′-deoxy-2′-fluoro; “m”: 2′-*O*-methyl; “P”: 5′-phosphate. Any substituents are named using standard chemical nomenclature.

**Table S3**. Panel addendum of unconjugated siRNAs for silencing efficacy screen and potency assays.

| Duplex ID | Target | Strand ID | Sequence (5′ to 3′) |
|---|---|---|---|
| *Positive Control* | | | |
| **D24** | *HTT* | **G1** | P(mU)#(fU)#(mA)(fA)(mU)(fC)(mU)(fC)(mU)(fU)(mU)(fA)(mC)#(fU)#(mG)#(fA)#(mU)#(fA)#(mU)#(fA) |
| | | **P2** | (fC)#(mA)#(fG)(mU)(fA)(mA)(fA)(mG)(fA)(mG)(fA)(mU)(fU)#(mA)#(fA) |
| *Benchmark from Primary Panel* | | | |
| **D25** | *HTT* | **G2** | methyl-P(mU)#(fU)#(mA)(fA)(mU)(fC)(mU)(fC)(mU)(fU)(mU)(fA)(mC)#(fU)#(mG)#(fA)#(mU)#(fA)#(mU)#(fA) |
| | | **P2** | (fC)#(mA)#(fG)(mU)(fA)(mA)(fA)(mG)(fA)(mG)(fA)(mU)(fU)#(mA)#(fA) |
| *Test Articles (Panel Addendum)* | | | |
| **D30** | *HTT* | **G25** | 2-methoxyethyl-P(mU)#(fU)#(mA)(fA)(mU)(fC)(mU)(fC)(mU)(fU)(mU)(fA)(mC)#(fU)#(mG)#(fA)#(mU)#(fA)#(mU)#(fA) |
| | | **P2** | (fC)#(mA)#(fG)(mU)(fA)(mA)(fA)(mG)(fA)(mG)(fA)(mU)(fU)#(mA)#(fA) |
| **D31** | *HTT* | **G26** | 2-(2-methoxyethoxy)ethyl-P(mU)#(fU)#(mA)(fA)(mU)(fC)(mU)(fC)(mU)(fU)(mU)(fA)(mC)#(fU)#(mG)#(fA)#(mU)#(fA)#(mU)#(fA) |
| | | **P2** | (fC)#(mA)#(fG)(mU)(fA)(mA)(fA)(mG)(fA)(mG)(fA)(mU)(fU)#(mA)#(fA) |
| **D32** | *HTT* | **G27** | isobutyl-P(mU)#(fU)#(mA)(fA)(mU)(fC)(mU)(fC)(mU)(fU)(mU)(fA)(mC)#(fU)#(mG)#(fA)#(mU)#(fA)#(mU)#(fA) |
| | | **P2** | (fC)#(mA)#(fG)(mU)(fA)(mA)(fA)(mG)(fA)(mG)(fA)(mU)(fU)#(mA)#(fA) |
| **D33** | *HTT* | **G28** | 4-pentenyl-P(mU)#(fU)#(mA)(fA)(mU)(fC)(mU)(fC)(mU)(fU)(mU)(fA)(mC)#(fU)#(mG)#(fA)#(mU)#(fA)#(mU)#(fA) |
| | | **P2** | (fC)#(mA)#(fG)(mU)(fA)(mA)(fA)(mG)(fA)(mG)(fA)(mU)(fU)#(mA)#(fA) |
| **D34** | *HTT* | **G29** | 4-pentynyl-P(mU)#(fU)#(mA)(fA)(mU)(fC)(mU)(fC)(mU)(fU)(mU)(fA)(mC)#(fU)#(mG)#(fA)#(mU)#(fA)#(mU)#(fA) |
| | | **P2** | (fC)#(mA)#(fG)(mU)(fA)(mA)(fA)(mG)(fA)(mG)(fA)(mU)(fU)#(mA)#(fA) |
| **D35** | *HTT* | **G30** | 2-butynyl-P(mU)#(fU)#(mA)(fA)(mU)(fC)(mU)(fC)(mU)(fU)(mU)(fA)(mC)#(fU)#(mG)#(fA)#(mU)#(fA)#(mU)#(fA) |
| | | **P2** | (fC)#(mA)#(fG)(mU)(fA)(mA)(fA)(mG)(fA)(mG)(fA)(mU)(fU)#(mA)#(fA) |

| | | | |
|---|---|---|---|
| **D36** | *HTT* | **G31** | 3-(1-naphthyl)propargyl-P(mU)#(fU)#(mA)(fA)(mU)(fC)(mU)(fC)(mU)(fU)(mU)(fA)(mC)#(fU)#(mG)#(fA)#(mU)#(fA)#(mU)#(fA) |
| | | **P2** | (fC)#(mA)#(fG)(mU)(fA)(mA)(fA)(mG)(fA)(mG)(fA)(mU)(fU)#(mA)#(fA) |
| **D37** | *HTT* | **G32** | 3-(3-pyridyl)propargyl-P(mU)#(fU)#(mA)(fA)(mU)(fC)(mU)(fC)(mU)(fU)(mU)(fA)(mC)#(fU)#(mG)#(fA)#(mU)#(fA)#(mU)#(fA) |
| | | **P2** | (fC)#(mA)#(fG)(mU)(fA)(mA)(fA)(mG)(fA)(mG)(fA)(mU)(fU)#(mA)#(fA) |
| **D38** | *HTT* | **G33** | 3-(4-pyridyl)propargyl-P(mU)#(fU)#(mA)(fA)(mU)(fC)(mU)(fC)(mU)(fU)(mU)(fA)(mC)#(fU)#(mG)#(fA)#(mU)#(fA)#(mU)#(fA) |
| | | **P2** | (fC)#(mA)#(fG)(mU)(fA)(mA)(fA)(mG)(fA)(mG)(fA)(mU)(fU)#(mA)#(fA) |
| **D39** | *HTT* | **G34** | 3-(2-thienyl)propargyl-P(mU)#(fU)#(mA)(fA)(mU)(fC)(mU)(fC)(mU)(fU)(mU)(fA)(mC)#(fU)#(mG)#(fA)#(mU)#(fA)#(mU)#(fA) |
| | | **P2** | (fC)#(mA)#(fG)(mU)(fA)(mA)(fA)(mG)(fA)(mG)(fA)(mU)(fU)#(mA)#(fA) |
| **D40** | *HTT* | **G35** | 3-aminopropyl-P(mU)#(fU)#(mA)(fA)(mU)(fC)(mU)(fC)(mU)(fU)(mU)(fA)(mC)#(fU)#(mG)#(fA)#(mU)#(fA)#(mU)#(fA) |
| | | **P2** | (fC)#(mA)#(fG)(mU)(fA)(mA)(fA)(mG)(fA)(mG)(fA)(mU)(fU)#(mA)#(fA) |
| **D41** | *HTT* | **G36** | 3-acetamidopropyl-P(mU)#(fU)#(mA)(fA)(mU)(fC)(mU)(fC)(mU)(fU)(mU)(fA)(mC)#(fU)#(mG)#(fA)#(mU)#(fA)#(mU)#(fA) |
| | | **P2** | (fC)#(mA)#(fG)(mU)(fA)(mA)(fA)(mG)(fA)(mG)(fA)(mU)(fU)#(mA)#(fA) |

Strand IDs are designated as follows: "G": guide strand; "P": passenger strand. Chemical modifications are designated as follows: "#": phosphorothioate linkage; "f": 2′-deoxy-2′-fluoro; "m": 2′-*O*-methyl; "P": 5′-phosphate. Any substituents are named using standard chemical nomenclature.

**Table S4**. Unconjugated siRNAs featuring additional 5′-P mimics for silencing potency assay.

| Duplex ID | Target | Strand ID | Sequence (5′ to 3′) |
|---|---|---|---|
| *Positive Control* | | | |
| **D24** | *HTT* | **G1** | P(mU)#(fU)#(mA)(fA)(mU)(fC)(mU)(fC)(mU)(fU)(mU)(fA)(mC)#(fU)#(mG)#(fA)#(mU)#(fA)#(mU)#(fA) |
| | | **P2** | (fC)#(mA)#(fG)(mU)(fA)(mA)(fA)(mG)(fA)(mG)(fA)(mU)(fU)#(mA)#(fA) |
| *Test Articles (Additional 5'-P Mimics)* | | | |
| **D42** | *HTT* | **G37** | PS(mU)#(fU)#(mA)(fA)(mU)(fC)(mU)(fC)(mU)(fU)(mU)(fA)(mC)#(fU)#(mG)#(fA)#(mU)#(fA)#(mU)#(fA) |
| | | **P2** | (fC)#(mA)#(fG)(mU)(fA)(mA)(fA)(mG)(fA)(mG)(fA)(mU)(fU)#(mA)#(fA) |
| **D43** | *HTT* | **G38** | phenylpropargyl-PS(mU)#(fU)#(mA)(fA)(mU)(fC)(mU)(fC)(mU)(fU)(mU)(fA)(mC)#(fU)#(mG)#(fA)#(mU)#(fA)#(mU)#(fA) |
| | | **P2** | (fC)#(mA)#(fG)(mU)(fA)(mA)(fA)(mG)(fA)(mG)(fA)(mU)(fU)#(mA)#(fA) |
| **D44** | *HTT* | **G39** | MsPA(mU)#(fU)#(mA)(fA)(mU)(fC)(mU)(fC)(mU)(fU)(mU)(fA)(mC)#(fU)#(mG)#(fA)#(mU)#(fA)#(mU)#(fA) |
| | | **P2** | (fC)#(mA)#(fG)(mU)(fA)(mA)(fA)(mG)(fA)(mG)(fA)(mU)(fU)#(mA)#(fA) |
| *Structural Comparandum (**G38** Analog)* | | | |
| **D28** | *HTT* | **G11** | phenylpropargyl-P(mU)#(fU)#(mA)(fA)(mU)(fC)(mU)(fC)(mU)(fU)(mU)(fA)(mC)#(fU)#(mG)#(fA)#(mU)#(fA)#(mU)#(fA) |
| | | **P2** | (fC)#(mA)#(fG)(mU)(fA)(mA)(fA)(mG)(fA)(mG)(fA)(mU)(fU)#(mA)#(fA) |

Strands are designated as follows: "G": guide strand; "P": passenger strand. Chemical modifications are designated as follows. "#": phosphorothioate linkage; "f": 2′-deoxy-2′-fluoro; "m": 2′-*O*-methyl; "P": 5′-phosphate; "PS": 5′-phosphorothioate; "MsPA"; 5′-mesylphosphoramidate. Any substituents are named using standard chemical nomenclature.

**Table S5**. Unconjugated siRNAs for silencing potency assay of second gene target (*MECP2*).

| Duplex ID | Target | Strand ID | Sequence (5′ to 3′) |
|---|---|---|---|
| *Positive Control* | | | |
| **D45** | *MECP2* | **G40** | P(mU)#(fA)#(mU)(fC)(mG)(fG)(mG)(fA)(mA)(fG)(mC)(fU)(mU)#(fU)#(mG)#(fU)#(mC)#(fA)#(mG)#(fA) |
| | | **P3** | (fC)#(mA)#(fA)(mA)(fG)(mC)(fU)(mU)(fC)(mC)(fC)(mG)(fA)#(mU)#(fA) |

| *Test Articles (Lead 5'-POR Variants)* | | | |
|---|---|---|---|
| **D46** | *MECP2* | **G41** | methyl-(mU)#(fA)#(mU)(fC)(mG)(fG)(mG)(fA)(mA)(fG)(mC)(fU)(mU)#(fU)#(mG)#(fU)#(mC)#(fA)#(mG)#(fA) |
| | | **P3** | (fC)#(mA)#(fA)(mA)(fG)(mC)(fU)(mU)(fC)(mC)(fC)(mG)(fA)#(mU)#(fA) |
| **D47** | *MECP2* | **G42** | phenylpropargyl-P(mU)#(fA)#(mU)(fC)(mG)(fG)(mG)(fA)(mA)(fG)(mC)(fU)(mU)#(fU)#(mG)#(fU)#(mC)#(fA)#(mG)#(fA) |
| | | **P3** | (fC)#(mA)#(fA)(mA)(fG)(mC)(fU)(mU)(fC)(mC)(fC)(mG)(fA)#(mU)#(fA) |

Strand IDs are designated as follows: "G": guide strand; "P": passenger strand. Chemical modifications are designated as follows: "#": phosphorothioate linkage; "f": 2′-deoxy-2′-fluoro; "m": 2′-*O*-methyl; "P": 5′-phosphate. Any substituents are named using standard chemical nomenclature.

***Silencing efficacy screen***. To screen mRNA silencing efficacy by passive uptake of cholesterol-conjugated siRNAs (**Table S1**), HeLa cells suspended in DMEM with 6% FBS were seeded into 96-well culture plates (10,000 cells/50 µL/well), followed by 3 µM siRNA (50 µL/well) in Opti-MEM (Gibco) for a final concentration of 1.5 µM siRNA in 3% FBS DMEM. Cells were incubated for 72 h at 37 °C and 5% $CO_2$, then processed (see *Cellular mRNA quantitation*).

To screen mRNA silencing efficacy by forward transfection of unconjugated siRNAs (**Table S3**), Lipofectamine RNAiMAX reagent (Invitrogen) was diluted in Opti-MEM (Gibco) at 1:50 v/v, then combined with an equal volume of 40 nM siRNA in Opti-MEM and incubated for 15 min to form 20 nM siRNA lipoplex. HeLa cells suspended in DMEM with 6% FBS were seeded into 96-well culture plates (10,000 cells/50 µL/well). siRNA lipoplexes were transferred to the culture plate (50 µL/well) for a final concentration of 10 nM siRNA lipoplex in 3% FBS DMEM. Cells were incubated for 72 h at 37 °C and 5% $CO_2$, then processed (see below).

***Silencing potency assay***. Using forward transfection of siRNAs (**Table S2**–**S5**), mRNA silencing potency was determined by dose–response assays. Lipofectamine RNAiMAX reagent (Invitrogen) was diluted in Opti-MEM (Gibco) at 1:50 v/v, then combined with an equal volume of 40 nM siRNA in Opti-MEM and incubated for 15 min to form 20 nM siRNA lipoplex. HeLa cells suspended in DMEM with 6% FBS were seeded into 96-well culture plates (10,000 cells/50 µL/well). In a deep-well plate, siRNA lipoplexes were diluted columnwise in Opti-MEM in a fivefold seven-point series (max. 20 nM), then transferred to the culture plate (50 µL/well) for a final dilution series of max. 10 nM siRNA lipoplex in 3% FBS DMEM. Cells were incubated for 72 h at 37 °C and 5% $CO_2$, then processed (see below).

***Cellular mRNA quantitation***. Target mRNA (*HTT* or *MECP2*) in the above assays was quantified and normalized to housekeeping mRNA (*HPRT*) using the QuantiGene 2.0 Singleplex (bDNA) assay kit (ThermoFisher) and validated probesets for *HTT* (SA-50339), *MECP2* (SA-11645), or *HPRT* (SB-15463). Cells were lysed in the 96-well culture plate with working QuantiGene Lysis Mixture (ThermoFisher; 250 µL/well) containing Proteinase K (ThermoFisher; 0.17 mg/mL), then digested at 55 °C for 30 min. Homogenized lysate aliquots (40 µL/well/probeset) were transferred to capture plate wells in triplicate. Target hybridization and signal amplification steps were performed per manufacturer protocol. Plate luminometry was conducted with a Tecan Spark microplate reader. Following subtraction of the mean blank signal, target mRNA absolute signal was normalized to *HPRT* mRNA absolute signal; target mRNA expression was calculated relative to mean untreated sample signal.

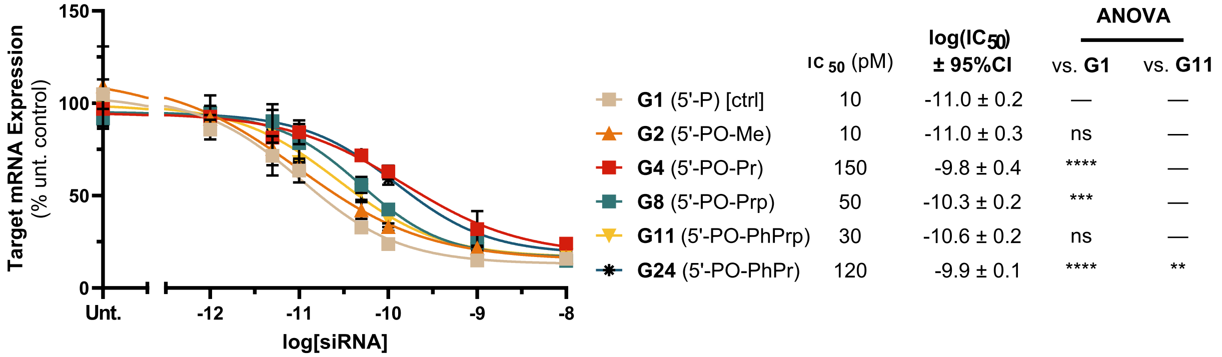


**Figure S3.** Dose–response assay of 5′-POR hits (**G2**, **G4, G8, G11**), a **G11** analog (**G24**), and control (**G1**) targeting *HTT* (**Table S2**). Potency and ANOVA of select pairwise comparisons shown (Šidák correction; $\alpha = 0.05$; * $p < 0.05$, ** $p < 0.01$, *** $p < 0.001$, **** $p < 0.0001$).

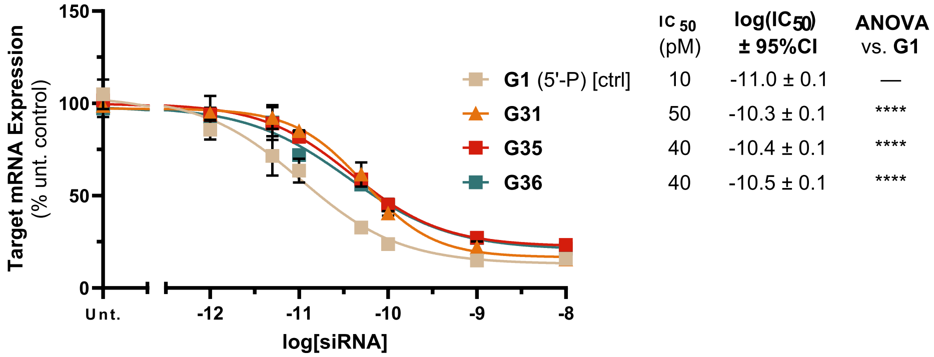


**Figure S4.** Dose–response assay of selection from 5′-POR panel addendum (**G31**, **G35, G36**) and control (**G1**) targeting *HTT* (**Table S3**). Potency and ANOVA of select pairwise comparisons shown (Šidák correction; $\alpha = 0.05$; * $p < 0.05$, ** $p < 0.01$, *** $p < 0.001$, **** $p < 0.0001$).

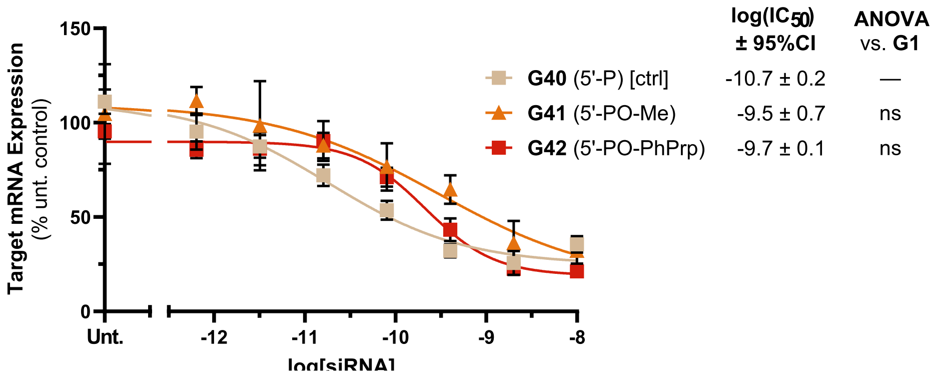


**Figure S5.** Dose–response assay of 5′-POR leads (**G41**, **G42**) and control (**G40**) targeting *MECP2* (**Table S5**). ANOVA of select pairwise comparisons shown Potency and ANOVA of select pairwise comparisons shown (Šidák correction; $\alpha = 0.05$; * $p < 0.05$, ** $p < 0.01$, *** $p < 0.001$, **** $p < 0.0001$).

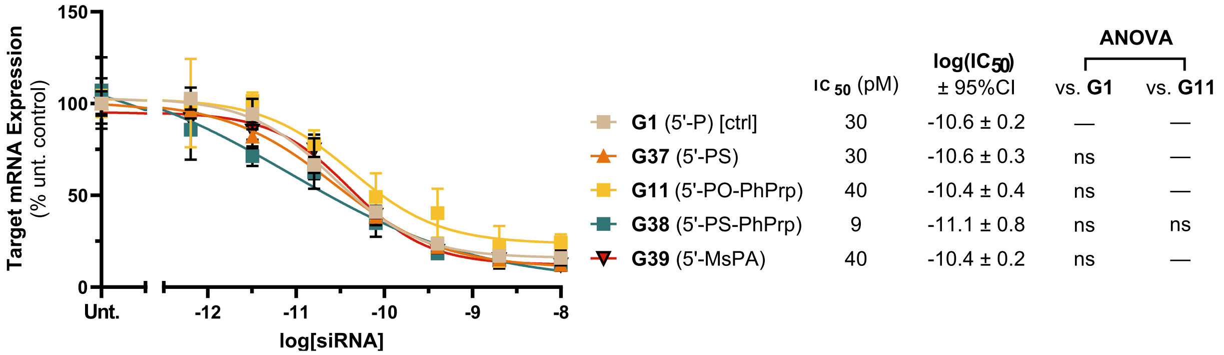


**Figure S6.** Dose–response assay of additional 5′-P mimics (**G37**–**G39**), a **G38** analog (**G11**), and control (**G1**) targeting *HTT* (**Table S4**). Potency and ANOVA of select pairwise comparisons shown (Šidák correction; α = 0.05; * p < 0.05, ** p < 0.01, *** p < 0.001, **** p < 0.0001).

### In Vitro Enzyme Susceptibility Assays

**Table S6**. Materials for phosphatase susceptibility assay (HPLC).

| Strand ID | Sequence (5′ to 3′) | $t_R$ (min) |
|---|---|---|
| *Test Articles/Standards* | | |
| **G1** | P(mU)#(fU)#(mA)(fA)(mU)(fC)(mU)(fC)(mU)(fU)(mU)(fA)(mC)#(fU)#(mG)#(fA)#(mU)#(fA)#(mU)#(fA) | 20.3 |
| **G2** | methyl-P(mU)#(fU)#(mA)(fA)(mU)(fC)(mU)(fC)(mU)(fU)(mU)(fA)(mC)#(fU)#(mG)#(fA)#(mU)#(fA)#(mU)#(fA) | 20.8 |
| **G11** | phenylpropargyl-P(mU)#(fU)#(mA)(fA)(mU)(fC)(mU)(fC)(mU)(fU)(mU)(fA)(mC)#(fU)#(mG)#(fA)#(mU)#(fA)#(mU)#(fA) | 23.9 |
| *Standard Only* | | |
| **G43** | (mU)#(fU)#(mA)(fA)(mU)(fC)(mU)(fC)(mU)(fU)(mU)(fA)(mC)#(fU)#(mG)#(fA)#(mU)#(fA)#(mU)#(fA) | 21.2 |

Chemical modifications are designated as follows. "#": phosphorothioate linkage; "f": 2′-deoxy-2′-fluoro; "m": 2′-*O*-methyl; "P": 5′-phosphate. Any substituents are named using standard chemical nomenclature. Retention times ($t_R$) used for analyte identification shown.

**Table S7**. Test articles/standards for phosphatase susceptibility assay (LC-MS)

| Strand ID | Sequence (5′ to 3′) |
|---|---|
| **G1** | P(mU)#(fU)#(mA)(fA)(mU)(fC)(mU)(fC)(mU)(fU)(mU)(fA)(mC)#(fU)#(mG)#(fA)#(mU)#(fA)#(mU)#(fA) |
| **G2** | methyl-P(mU)#(fU)#(mA)(fA)(mU)(fC)(mU)(fC)(mU)(fU)(mU)(fA)(mC)#(fU)#(mG)#(fA)#(mU)#(fA)#(mU)#(fA) |
| **G11** | phenylpropargyl(mU)#(fU)#(mA)(fA)(mU)(fC)(mU)(fC)(mU)(fU)(mU)(fA)(mC)#(fU)#(mG)#(fA)#(mU)#(fA)#(mU)#(fA) |
| **G37** | PS(mU)#(fU)#(mA)(fA)(mU)(fC)(mU)(fC)(mU)(fU)(mU)(fA)(mC)#(fU)#(mG)#(fA)#(mU)#(fA)#(mU)#(fA) |
| **G38** | phenylpropargyl-PS(mU)#(fU)#(mA)(fA)(mU)(fC)(mU)(fC)(mU)(fU)(mU)(fA)(mC)#(fU)#(mG)#(fA)#(mU)#(fA)#(mU)#(fA) |
| **G43** | (mU)#(fU)#(mA)(fA)(mU)(fC)(mU)(fC)(mU)(fU)(mU)(fA)(mC)#(fU)#(mG)#(fA)#(mU)#(fA)#(mU)#(fA) |

Chemical modifications are designated as follows: "#": phosphorothioate linkage; "f": 2′-deoxy-2′-fluoro; "m": 2′-*O*-methyl; "P": 5′-phosphate; "PS": 5′-phosphorothioate; "MsPA"; 5′-mesylphosphoramidate. Any substituents are named using standard chemical nomenclature. Calculated *m*/*z* used for analyte identification included in **Table S11**.

**Table S8**. Materials for 5′-exonuclease susceptibility assay

| Strand ID | Sequence (5′ to 3′) | $t_R$ (min) |
|---|---|---|
| *Test Articles/Standards* | | |
| **ON1** | P(mU)(fU)(mA)(fA)(mU)(fC)(mU)(fC)(mU)(fU)(mU)(fA)(mC)(fU)(mG)(fA)(mU)(fA)(mU)(fA) | 14.2 |

| | | |
|---|---|---|
| **ON2** | (mU)(fU)(mA)(fA)(mU)(fC)(mU)(fC)(mU)(fU)(mU)(fA)(mC)(fU)(mG)(fA)(mU)(fA)(mU)(fA) | 14.3 |
| **ON3** | VP(mU)(fU)(mA)(fA)(mU)(fC)(mU)(fC)(mU)(fU)(mU)(fA)(mC)(fU)(mG)(fA)(mU)(fA)(mU)(fA) | 14.3 |
| **ON4** | methyl-P(mU)(fU)(mA)(fA)(mU)(fC)(mU)(fC)(mU)(fU)(mU)(fA)(mC)(fU)(mG)(fA)(mU)(fA)(mU)(fA) | 14.3 |
| **ON5** | phenylpropargyl-P(mU)(fU)(mA)(fA)(mU)(fC)(mU)(fC)(mU)(fU)(mU)(fA)(mC)(fU)(mG)(fA)(mU)(fA)(mU)(fA) | 15.5 |
| **ON6** | PS(mU)(fU)(mA)(fA)(mU)(fC)(mU)(fC)(mU)(fU)(mU)(fA)(mC)(fU)(mG)(fA)(mU)(fA)(mU)(fA) | 14.2 |
| **ON7** | phenylpropargyl-PS(mU)(fU)(mA)(fA)(mU)(fC)(mU)(fC)(mU)(fU)(mU)(fA)(mC)(fU)(mG)(fA)(mU)(fA)(mU)(fA) | 16.6 |
| **ON8** | MsPA(mU)(fU)(mA)(fA)(mU)(fC)(mU)(fC)(mU)(fU)(mU)(fA)(mC)(fU)(mG)(fA)(mU)(fA)(mU)(fA) | 15.0 |
| *Internal Standard* | | |
| **ON9** | (dT)(dT)(dT)(dT)(dT)(dT)(dT)(dT) | 9.1 |

Chemical modifications are designated as follows. "f": 2′-deoxy-2′-fluoro; "m": 2′-*O*-methyl; "d": 2′-deoxy; "P": 5′-phosphate; "PS": 5′-phosphorothioate; "VP": 5′-(*E*)-vinylphosphonate. Any substituents are named using standard chemical nomenclature. Retention times ($t_R$) used for identification of intact analytes and internal standard shown.

***Phosphatase susceptibility assays***. For the short-duration HPLC assay, test articles (**Table S6**; 1.5 μM, 750 pmol) were incubated with Quick CIP (New England Biolabs, Lot 10187347; 25 μU/mL) in rCutSmart buffer (New England Biolabs) at 37 °C. From the reaction mixtures, samples (100 μL, 150 pmol) were collected directly after mixing (~0 min) and at 5, 15, 30, and 60 min. All samples were immediately denatured at 95 °C for 5 min, then flash-frozen in liquid nitrogen. HPLC analysis was performed on an Agilent 1260 Infinity instrument using the following conditions: Hamilton PRP-C18 column, 60 °C, 1 mL/min; buffer A: 100 mM hexafluoroisopropanol, 15 mM $Et_3N$ in $H_2O$; buffer B: 100 mM hexafluoroisopropanol, 15 mM $Et_3N$ in MeCN; gradient profile: 5–30% B for 30 min, 100% B for 2.5 min, 5% B for 2.5 min; $A_{260}$ peak monitoring. Standards were analyzed to establish retention times (see **Table S3**); co-injections of **G43** with the other standards were also analyzed to validate peak resolution. Intact analyte and 5′-OH degradant in each sample were quantified by integrating the respective $A_{260}$ peaks in LC OpenLab ChemStation (Agilent), processing the data in Microsoft Excel, and plotting in GraphPad Prism (**Fig. S7**).

For the long-duration LC-MS assay, test articles (**Table S7**; 10 μM, 2 nmol) were incubated with Quick CIP (New England Biolabs, Lot 10226662; 0.1 mU/mL) in rCutSmart buffer (New England Biolabs) at 37 °C. From the reaction mixtures, samples (20 μL, 200 pmol) were collected directly after mixing (~0 min) and at 5 min, 10 min, 20 min, 1 h, 3 h, 5 h, 8 h, and 12 h. All samples were immediately quenched with 6.25 mM EDTA in $H_2O$ (80 μL), denatured at 95 °C for 5 min, then flash-frozen in liquid nitrogen. Sample composition was characterized by standard LC-MS analysis (see *Oligonucleotide Characterization and Purification*).

***5′-Exonuclease susceptibility assay***. Test articles (**Table S8**; 10 μM, 2.4 nmol) were incubated with internal standard (**ON9**; 10 μM, 2.4 nmol) and XRN1 (New England Biolabs, Lot 10274120; 30 U/mL) in NEBuffer 3 (New England Biolabs) at 30 °C. From the reaction mixtures, samples (20 μL, 200 pmol) were collected directly after mixing (~0 min) and at 10 min, 20 min, 40 min, 1 h, 1.5 h, 3 h, 5 h, 8 h, and 12 h (0 min and 12 h samples in duplicate). All samples were immediately quenched with 6.25 mM EDTA in $H_2O$ (80 μL), denatured at 95 °C for 5 min, then flash-frozen in liquid nitrogen. HPLC analysis was performed on an Agilent 1260 Infinity instrument using the following conditions: Agilent PL-SAX column, 50 °C, 1 mL/min; buffer A: 25 mM $NaH_2PO_4$ in $H_2O$, pH 7; buffer B: 1 M $NaClO_4$, 25 mM $NaH_2PO_4$ in $H_2O$, pH 7); gradient profile: 5–40% B for 20 min, 100% B for 2.5 min, 5% B for 2.5 min; $A_{260}$ peak monitoring. Standards were analyzed to establish retention times (**Table S8**). Intact test article and **ON9** in each sample were quantified by integrating the respective $A_{260}$ peaks in LC OpenLab ChemStation (Agilent); test article amount was normalized to internal standard and plotted in GraphPad Prism. The duplicate samples for 0 min and 12 h were characterized by standard LC-MS analysis (see *Oligonucleotide Characterization and Purification*).

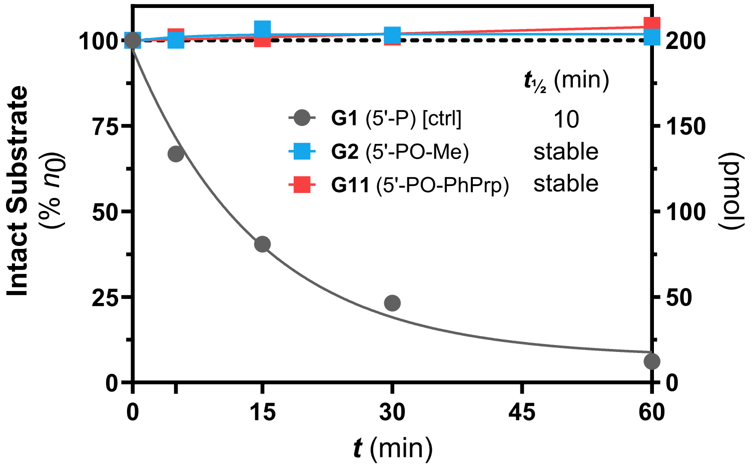


**Figure S7.** Phosphatase susceptibility of lead 5′-POR and 5′-P control guide strands (Table over 1 h (HPLC analysis).

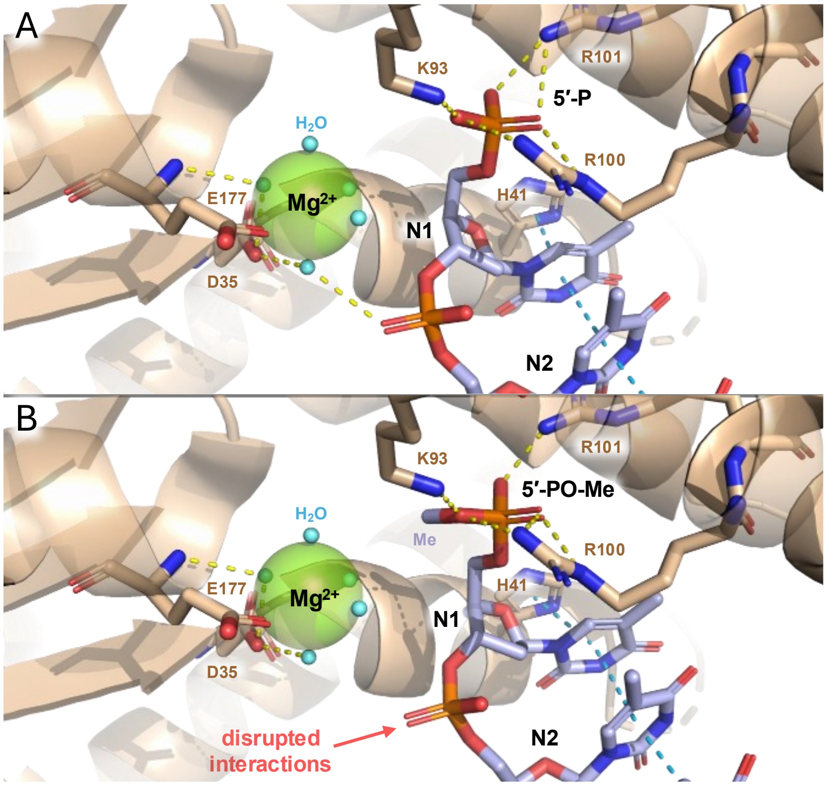


**Figure S8. (A)** XRN1 with bound 5′-P ssDNA (PDBID: 2Y35). **(B)** XRN1 with 5′-PO-Me ssDNA model, showing disrupted interactions between the magnesium inner hydration shell and the first internucleotide linkage. Note the magnesium ion coordinated to D35 and five waters in the catalytic center, extensive salt bridge network in the 5′-P binding pocket (K93, R100, R101), and π-π stack formed by H41 and the N1, N2 nucleobases. Salt bridges/hydrogen bonds (dashed yellow) and π-stacking (dashed blue) predicted in PyMOL.

### In Cellula Metabolism and RISC Loading Studies

***Lysate preparation***. HeLa cells were grown to 80% confluency on 15 cm culture plates (see *Cell culture*), washed gently with phosphate-buffered saline (PBS; Gibco), then lipofected with siRNA (**D28**; see **Table S2**) as follows. Lipofectamine RNAiMAX reagent (Invitrogen; 50 µL/plate) was diluted in Opti-MEM (Gibco; 2.5 mL/plate), then combined with siRNA (300 pmol/plate) in Opti-MEM (2.5 mL/plate) and incubated for 15 min to form siRNA lipoplex (300 pmol/5 mL/plate). The siRNA lipoplex was then combined with DMEM with 10% FBS (Gibco; 25 mL/plate) and transferred to each culture plate (30 mL/plate). The plates were incubated for 48 h at 37 °C and 5% $CO_2$, then cells were trypsinized, pelleted, incubated for 10 min on ice in cold NETN lysis buffer (50 mM Tris, 150 mM NaCl, 5 mM EDTA, 0.5% w/v NP-40, 10% w/v glycerol, 0.5 mM dithiothreitol, pH 7.5; 1 mL/plate) with cOmplete Protease Inhibitor Cocktail (Roche), and centrifuged (15,000 g, 10 min, 4 °C). The supernatant (debris-free lysate) was collected and either prepared for mass analysis or used for RISC pulldown (see below). For analytical samples, lysate was mixed 2:1 by volume with Quantigene Lysis Mixture (Invitrogen) containing Proteinase K (ThermoFisher; 0.5 mg/mL) and incubated at 65 °C for 30 min. The proteolyzed lysate was then mixed 1:2 by volume with Ampure XP bead slurry (Beckman

Coulter) and incubated at room temperature for 5 min. The supernatant was collected from immobilized beads, supplemented 1:0.25 by volume with borax buffer (150 mM borax, 150 mM boric acid) and 1:0.05 by volume with 2 M $MgCl_2$, shaken overnight at 65°C, and precipitated with cold *i*PrOH; precipitate was washed with 80% aq. EtOH then resuspended in nuclease-free $H_2O$.

***RISC pulldown***. Pierce Glutathione Magnetic Agarose Bead slurry (Invitrogen; 70 µL) was washed thrice with 0.02% w/v Tween-20 (Sigma-Aldrich) in cold PBS. Washed beads were incubated with purified rGST-T6B (140 µg) for 3 h at 4 °C; the supernatant was then discarded, and the beads were washed thrice with 0.02% w/v Tween-20 in cold PBS with 1 mM dithiothreitol (ThermoFisher) to remove unbound rGST-T6B. Lysate (input; ~1 mL) was added over the washed beads and incubated overnight at 4 °C to capture the lysate RISC pool. Subsequently, the beads were washed five times with cold NETN lysis buffer (see *Lysate preparation*) to remove all unbound matrix; before washing, an aliquot of the supernatant (~100 µL) was reserved for analysis. An aliquot of the resultant pulldown bead bed (corresponding to 1 culture plate) was also reserved for protein analysis (see below). For siRNA guide analysis (see below), the remaining pulldown beads were incubated in 10% v/v piperidine in $H_2O$ at 95 °C for 90 min to denature the captured RISC pool and digest endogenous RNA. The supernatant was collected, piperidine was removed by lyophilization, and the sample (pulldown fraction) was precipitated with cold *i*PrOH; precipitate was washed with 80% aq. EtOH then resuspended in nuclease-free $H_2O$.

***Western blot analysis.*** The pulldown fraction was prepared from the bead bed (corresponding to 1 culture plate) by incubating the beads in 50 µL of working NuPAGE LDS sample buffer (Invitrogen) at 95 °C for 10 min, then flash-cooling on ice. Aliquots of lysate input and supernatant (5 µL each) were also incubated in working sample buffer. Samples were separated via SDS-PAGE on a NuPAGE 4–12% Bis-Tris gel (Invitrogen) with a Spectra broad-range protein ladder (Invitrogen). The gel was transferred to an Immobilon-FL PVDF membrane (Milipore) with a Trans-Blot Turbo semi-dry transfer system (Bio-Rad) and blocked with Intercept PBS blocking buffer (LICORbio) for 1 h at room temperature. For immunodetection of AGO2 (analyte) and GAPDH (control), the membrane was incubated at 4 °C overnight with the following primary antibody dilutions in blocking buffer: rabbit anti–Argonaute-2 IgG [EPR10410] (Abcam, Lot 1029045-19, 1:1,000) and rabbit anti–GAPDH [14C10] IgG (Cell Signaling, Lot 16, 1:40,000). After washing thrice in PBST buffer, the membrane was incubated with goat anti-rabbit IgG H+L secondary antibody with AlexaFluor Plus 800 conjugate (Invitrogen, Lot 3214509, 1:20,000) for 1 h at room temperature. The membrane was again washed thrice with PBST buffer, visualized using a Gel Doc Go fluorescence gel imager (Bio-Rad), and analyzed with ImageJ software (**Fig. S9**).

***Guide analysis***. Analytical lysate samples and **G1**, **G11** standards (100 pmol) were characterized by LC-MS analysis on an Agilent 6530 Accurate Mass Q-TOF LC-MS instrument; pulldown fractions and **G1**, **G11** standards (1 pmol) were characterized by LC-MS analysis on an Agilent

6230 LC/TOF instrument. LC conditions were as follows: AdvanceBio Oligonucleotide C18 LC column (Agilent), 60 °C, 0.85 mL/min; buffer A: 50 mM hexafluoroisopropanol, 10 mM $Et_3N$ in $H_2O$; buffer B: 50 mM hexafluoroisopropanol, 10 mM $Et_3N$ in MeOH; gradient profile: 5% B for 0.7 min, 5–100% B for 5.5 min, 5% B for 1.5 min (lysate samples) or 5% B for 1.6 min, 5–100% B for 5.5 min, 100% B for 1.6 min, 5% B for 1.6 min (pulldown samples); $A_{260}$ peak monitoring. ESI-MS parameters were as follows: negative-ion profile mode, 4,000 V capillary potential, 100–3,200 m/z range, 2 Hz scan rate. Mass spectra were extracted in Agilent BioConfirm and plotted in GraphPad Prism.

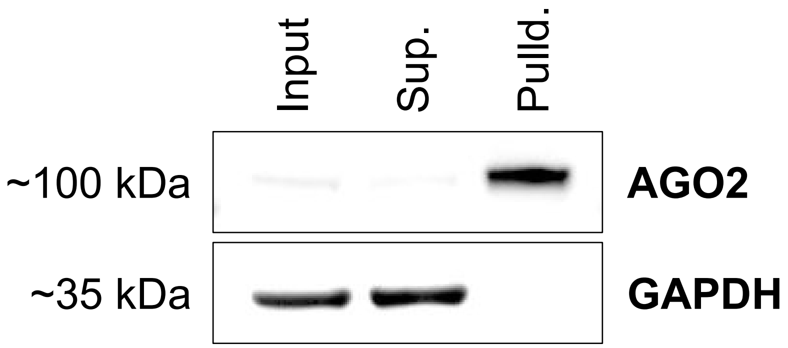


**Figure S9.** Western blot of **D28** lysate (input), supernatant, and pulldown fraction for human Argonaute-2 analyte (AGO2, MW ~97 kDa) and glyceraldehyde-3-phosphate dehydrogenase control (GAPDH, MW ~36 kDa). GAPDH (unbound control) is depleted, whereas AGO2 (binding target) is strongly enriched.

### Biochemical and Structural Analyses

**Table S9**. Oligonucleotides for AGO2 co-crystallization

| Strand ID | Sequence (5′ to 3′) |
|---|---|
| **ON10** | phenylpropargyl-P(mU)#(fG)#(mG)(fA)(fG)(fU)(mG)(fU)(mG)(fA)(mC)(fA)(mA)(fU)(mG)(fG)(mU)(mG)(mU)#(mU)#(fU) |
| **ON11** | 3-phenylpropyl-P(mU)#(fG)#(mG)(fA)(fG)(fU)(mG)(fU)(mG)(fA)(mC)(fA)(mA)(fU)(mG)(fG)(mU)(mG)(mU)#(mU)#(fU) |
| **ON12** | methyl-P(mU)#(fG)#(mG)(fA)(fG)(fU)(mG)(fU)(mG)(fA)(mC)(fA)(mA)(fU)(mG)(fG)(mU)(mG)(mU)#(mU)#(fU) |
| **ON13** | P(mU)#(fG)#(mG)(fA)(fG)(fU)(mG)(fU)(mG)(fA)(mC)(fA)(mA)(fU)(mG)(fG)(mU)(mG)(mU)#(mU)#(fU) |

Chemical modifications are designated as follows. "#": phosphorothioate linkage; "f": 2′-deoxy-2′-fluoro; "m": 2′-*O*-methyl; "P": 5′-phosphate. Any substituents are named using standard chemical nomenclature.

**Table S10.** Data collection and refinement statistics.

| | AGO2:ON10 | AGO2:ON11 |
|---|---|---|
| PDBID | 9OBD | 9OBE |
| Wavelength | 0.976 | 1 |
| Resolution range | 64.68 - 2.02 (2.04 - 2.02) | 60.17 - 2.3 (2.38 - 2.3) |
| Space group | P 1 21 1 | P 1 21 1 |
| Unit cell | 62.905 108.926 67.929 90 107.8 90 | 63.116 108.236 68.529 90 107.59 90 |
| Total reflections | 793562 (26550) | 265112 (27133) |
| Unique reflections | 56596 (1828) | 38542 (3846) |
| Multiplicity | 14.0 (14.5) | 6.9 (7.1) |
| Completeness (%) | 98.67 (94.66) | 98.50 (99.37) |
| Mean I/sigma(I) | 6.85 (0.63) | 17.75 (4.10) |
| Wilson B-factor | 35.62 | 46.22 |

| | | |
|---|---|---|
| R-merge | 0.2738 (4.428) | 0.05461 (0.4051) |
| R-meas | 0.2841 (4.588) | 0.05908 (0.4379) |
| R-pim | 0.07514 (1.192) | 0.02229 (0.1645) |
| CC1/2 | 0.996 (0.291) | 0.999 (0.946) |
| CC* | 0.999 (0.671) | 1 (0.986) |
| Reflections used in refinement | 56572 (1824) | 38521 (3843) |
| Reflections used for R-free | 2809 (90) | 1915 (195) |
| R-work | 0.2317 (0.3802) | 0.2073 (0.2813) |
| R-free | 0.2670 (0.3524) | 0.2539 (0.3123) |
| Number of non-hydrogen atoms | 7121 | 6876 |
| macromolecules | 6554 | 6495 |
| ligands | 257 | 220 |
| solvent | 310 | 161 |
| Protein residues | 810 | 803 |
| RMS(bonds) | 0.020 | 0.024 |
| RMS(angles) | 1.02 | 1.07 |
| Ramachandran favored (%) | 97.49 | 97.35 |
| Ramachandran allowed (%) | 2.51 | 2.65 |
| Ramachandran outliers (%) | 0.00 | 0.00 |
| Rotamer outliers (%) | 1.11 | 1.68 |
| Clashscore | 2.14 | 3.00 |
| Average B-factor | 45.74 | 57.26 |
| macromolecules | 45.81 | 57.30 |
| ligands | 45.35 | 55.84 |
| solvent | 43.21 | 48.41 |

Statistics for the highest-resolution shell are shown in parentheses.

***Preparation of recombinant human AGO2 loaded with modified oligonucleotides.*** Human AGO2 was purified according to a published method.[1] Briefly, human AGO2 was expressed in Sf9 cells using the baculovirus system. Cell pellets were lysed, and N-terminally His6-tagged AGO2 was purified by Ni-NTA affinity chromatography. Synthetic single-strand oligonucleotides (**ON10**–**ON13**) were each incubated with the isolated AGO2, and the His6-tag was removed using tobacco etch virus protease during overnight dialysis at 4 °C. AGO2 molecules loaded with the synthetic oligonucleotide were captured using an antisense oligonucleotide (IDT) and eluted with a biotinylated DNA competitor (IDT). The competitor was removed with neutravidin resin (Pierce) and by ion exchange chromatography with Q resin (Cytiva). Protein concentration was determined on Nanodrop OneC (Thermo Scientific) using a calculated extinction coefficient at 280 nm of 174120 M-1 cm-1 and a molecular mass of 108 kDa.

***Crystallization.*** The AGO2:siRNAs stocks were diluted to 2 mg/ml and crystallized by screening around a published condition[2] in 24-well plates (Hampton). Final crystallization conditions were 16% w/v PEG3350, 100 mM phenol, 12% v/v propan-2-ol, and 100 mM Tris-HCl (pH 8), yielding elongated hexagons. The crystals were prepared for freezing by transferring to a solution of 18% w/v PEG 3350, 100 mM Tris-HCl pH 8, and 25% w/v ethylene glycol as cryoprotectant.

***X-ray diffraction data collection and processing.*** Data was collected at ALS Beamline 5.0.2 and 5.0.3. Diffraction data were indexed and integrated using DIALS,[3] then scaled and merged in Aimless.[4] Initial phases were generated by molecular replacement using Phaser[5] with the protein component of Ago2 bound to an unrelated guide as search model (PDBID: 4W5N). The RNA was modelled in Coot.[6] Modified nucleotides were generated using eLBOW.[7] The model was iteratively improved with phenix.refine,[8] and manual adjustments in Coot.[6]

***Radiolabeling of target RNA.*** Synthetic RNA oligonucleotide full miR-122 target to 19 (FT19) (IDT) was radiolabeled at the 5′-end using γ-phosphate $^{32}$P-labeled ATP (Revvity Health Sciences Inc). Labeled RNAs were purified by denaturing polyacrylamide gel followed by gel extraction and ethanol precipitation. RNA concentration was determined by $A_{260}$ on a Nanodrop OneC (Thermo Scientific) using the extinction coefficient calculated by the IDT tool.

***AGO2 single-turnover cleavage kinetics.*** Reactions were performed in 30 mM Tris-HCl (pH 8.0), 0.1 M KOAc, 2 mM $Mg(OAc)_2$, 0.5 mM TCEP, with a total of 0.2 nM radiolabeled RNA, and 0.1 mg/ml *S. cerevisiae* tRNA. After removal of a 0 s time point, the reactions were started by addition of 5/10/20 nM AGO2:guide complex. Time points were taken by transferring 10 µl reaction to 10 µl 2× formamide loading buffer. The time points were analyzed by denaturing polyacrylamide gels (15% acrylamide, 8 M urea) in 0.5× TBE. A phosphor screen (Cytiva) was exposed for visualization, imaged on a Typhoon phosphorimager (Cytiva), and quantified using ImageQuant (Cytiva). Prism (GraphPad Software) was used to fit the data to a one-phase decay curve.

***Visualization.*** Structures were visualized in Pymol (Schrodinger) and ChimeraX (UCSF).[9]

### Oligonucleotide Authentication

**Table S11**. Mass analysis of all oligonucleotides in this work.

| Strand ID | Sequence (5′ to 3′) | Calc. *m/z* | Obs. *m/z* |
|---|---|---|---|
| **G1** | P(mU)#(fU)#(mA)(fA)(mU)(fC)(mU)(fC)(mU)(fU)(mU)(fA)(mC)#(fU)#(mG)#(fA)#(mU)#(fA)#(mU)#(fA) | 6620.44 | 6619.64 |
| **G2** | methyl-P(mU)#(fU)#(mA)(fA)(mU)(fC)(mU)(fC)(mU)(fU)(mU)(fA)(mC)#(fU)#(mG)#(fA)#(mU)#(fA)#(mU)#(fA) | 6556.49 | 6555.66 |
| **G3** | ethyl-P(mU)#(fU)#(mA)(fA)(mU)(fC)(mU)(fC)(mU)(fU)(mU)(fA)(mC)#(fU)#(mG)#(fA)#(mU)#(fA)#(mU)#(fA) | 6570.53 | 6570.04 |
| **G4** | propyl-P(mU)#(fU)#(mA)(fA)(mU)(fC)(mU)(fC)(mU)(fU)(mU)(fA)(mC)#(fU)#(mG)#(fA)#(mU)#(fA)#(mU)#(fA) | 6584.56 | 6583.77 |
| **G5** | butyl-P(mU)#(fU)#(mA)(fA)(mU)(fC)(mU)(fC)(mU)(fU)(mU)(fA)(mC)#(fU)#(mG)#(fA)#(mU)#(fA)#(mU)#(fA) | 6598.59 | 6598.00 |
| **G6** | pentyl-P(mU)#(fU)#(mA)(fA)(mU)(fC)(mU)(fC)(mU)(fU)(mU)(fA)(mC)#(fU)#(mG)#(fA)#(mU)#(fA)#(mU)#(fA) | 6612.61 | 6612.47 |
| **G7** | hexyl-P(mU)#(fU)#(mA)(fA)(mU)(fC)(mU)(fC)(mU)(fU)(mU)(fA)(mC)#(fU)#(mG)#(fA)#(mU)#(fA)#(mU)#(fA) | 6626.64 | 6626.66 |
| **G8** | propargyl-P(mU)#(fU)#(mA)(fA)(mU)(fC)(mU)(fC)(mU)(fU)(mU)(fA)(mC)#(fU)#(mG)#(fA)#(mU)#(fA)#(mU)#(fA) | 6580.53 | 6580.07 |
| **G9** | 3-butynyl-P(mU)#(fU)#(mA)(fA)(mU)(fC)(mU)(fC)(mU)(fU)(mU)(fA)(mC)#(fU)#(mG)#(fA)#(mU)#(fA)#(mU)#(fA) | 6594.55 | 6593.88 |
| **G10** | 5-hexynyl-P(mU)#(fU)#(mA)(fA)(mU)(fC)(mU)(fC)(mU)(fU)(mU)(fA)(mC)#(fU)#(mG)#(fA)#(mU)#(fA)#(mU)#(fA) | 6622.61 | 6621.88 |
| **G11** | 1-phenylpropargyl-P(mU)#(fU)#(mA)(fA)(mU)(fC)(mU)(fC)(mU)(fU)(mU)(fA)(mC)#(fU)#(mG)#(fA)#(mU)#(fA)#(mU)#(fA) | 6656.62 | 6656.26 |
| **G12** | 2-hydroxyethyl-P(mU)#(fU)#(mA)(fA)(mU)(fC)(mU)(fC)(mU)(fU)(mU)(fA)(mC)#(fU)#(mG)#(fA)#(mU)#(fA)#(mU)#(fA) | 6586.53 | 6585.98 |

| | | | |
|---|---|---|---|
| **G13** | 3-hydroxypropyl-P(mU)#(fU)#(mA)(fA)(mU)(fC)(mU)(fC)(mU)(fU)(mU)(fA)(mC)#(fU)#(mG)#(fA)#(mU)#(fA)#(mU)#(fA) | 6600.56 | 6599.99 |
| **G14** | 4-hydroxybutyl-P(mU)#(fU)#(mA)(fA)(mU)(fC)(mU)(fC)(mU)(fU)(mU)(fA)(mC)#(fU)#(mG)#(fA)#(mU)#(fA)#(mU)#(fA) | 6614.58 | 6613.46 |
| **G15** | 6-hydroxyhexyl-P(mU)#(fU)#(mA)(fA)(mU)(fC)(mU)(fC)(mU)(fU)(mU)(fA)(mC)#(fU)#(mG)#(fA)#(mU)#(fA)#(mU)#(fA) | 6642.64 | 6641.78 |
| **G16** | 2-phosphonooxyethyl-P(mU)#(fU)#(mA)(fA)(mU)(fC)(mU)(fC)(mU)(fU)(mU)(fA)(mC)#(fU)#(mG)#(fA)#(mU)#(fA)#(mU)#(fA) | 6666.50 | 6666.05 |
| **G17** | 3-phosphonooxypropyl-P(mU)#(fU)#(mA)(fA)(mU)(fC)(mU)(fC)(mU)(fU)(mU)(fA)(mC)#(fU)#(mG)#(fA)#(mU)#(fA)#(mU)#(fA) | 6680.53 | 6679.75 |
| **G18** | 4-phosphonooxybutyl-P(mU)#(fU)#(mA)(fA)(mU)(fC)(mU)(fC)(mU)(fU)(mU)(fA)(mC)#(fU)#(mG)#(fA)#(mU)#(fA)#(mU)#(fA) | 6694.56 | 6693.95 |
| **G19** | 6-phosphonooxyhexyl-P(mU)#(fU)#(mA)(fA)(mU)(fC)(mU)(fC)(mU)(fU)(mU)(fA)(mC)#(fU)#(mG)#(fA)#(mU)#(fA)#(mU)#(fA) | 6722.61 | 6721.56 |
| **G20** | allyl-P(mU)#(fU)#(mA)(fA)(mU)(fC)(mU)(fC)(mU)(fU)(mU)(fA)(mC)#(fU)#(mG)#(fA)#(mU)#(fA)#(mU)#(fA) | 6582.54 | 6581.95 |
| **G21** | benzyl-P(mU)#(fU)#(mA)(fA)(mU)(fC)(mU)(fC)(mU)(fU)(mU)(fA)(mC)#(fU)#(mG)#(fA)#(mU)#(fA)#(mU)#(fA) | 6632.60 | 6632.43 |
| **G22** | 2-phenylethyl-P(mU)#(fU)#(mA)(fA)(mU)(fC)(mU)(fC)(mU)(fU)(mU)(fA)(mC)#(fU)#(mG)#(fA)#(mU)#(fA)#(mU)#(fA) | 6646.63 | 6646.32 |
| **G23** | isopropyl-P(mU)#(fU)#(mA)(fA)(mU)(fC)(mU)(fC)(mU)(fU)(mU)(fA)(mC)#(fU)#(mG)#(fA)#(mU)#(fA)#(mU)#(fA) | 6584.56 | 6584.23 |
| **G24** | 3-phenylpropyl-P(mU)#(fU)#(mA)(fA)(mU)(fC)(mU)(fC)(mU)(fU)(mU)(fA)(mC)#(fU)#(mG)#(fA)#(mU)#(fA)#(mU)#(fA) | 6660.66 | 6660.09 |
| **G25** | 2-methoxyethyl-P(mU)#(fU)#(mA)(fA)(mU)(fC)(mU)(fC)(mU)(fU)(mU)(fA)(mC)#(fU)#(mG)#(fA)#(mU)#(fA)#(mU)#(fA) | 6600.56 | 6599.79 |
| **G26** | 2-(2-methoxyethoxy)ethyl-P(mU)#(fU)#(mA)(fA)(mU)(fC)(mU)(fC)(mU)(fU)(mU)(fA)(mC)#(fU)#(mG)#(fA)#(mU)#(fA)#(mU)#(fA) | 6644.61 | 6644.27 |
| **G27** | isobutyl-P(mU)#(fU)#(mA)(fA)(mU)(fC)(mU)(fC)(mU)(fU)(mU)(fA)(mC)#(fU)#(mG)#(fA)#(mU)#(fA)#(mU)#(fA) | 6598.48 | 6598.07 |
| **G28** | 4-pentenyl-P(mU)#(fU)#(mA)(fA)(mU)(fC)(mU)(fC)(mU)(fU)(mU)(fA)(mC)#(fU)#(mG)#(fA)#(mU)#(fA)#(mU)#(fA) | 6610.60 | 6609.77 |
| **G29** | 4-pentynyl-P(mU)#(fU)#(mA)(fA)(mU)(fC)(mU)(fC)(mU)(fU)(mU)(fA)(mC)#(fU)#(mG)#(fA)#(mU)#(fA)#(mU)#(fA) | 6608.58 | 6608.34 |
| **G30** | 2-butynyl-P(mU)#(fU)#(mA)(fA)(mU)(fC)(mU)(fC)(mU)(fU)(mU)(fA)(mC)#(fU)#(mG)#(fA)#(mU)#(fA)#(mU)#(fA) | 6594.55 | 6594.25 |
| **G31** | 3-(1-naphthyl)propargyl-P(mU)#(fU)#(mA)(fA)(mU)(fC)(mU)(fC)(mU)(fU)(mU)(fA)(mC)#(fU)#(mG)#(fA)#(mU)#(fA)#(mU)#(fA) | 6706.69 | 6705.66 |
| **G32** | 3-(3-pyridyl)propargyl-P(mU)#(fU)#(mA)(fA)(mU)(fC)(mU)(fC)(mU)(fU)(mU)(fA)(mC)#(fU)#(mG)#(fA)#(mU)#(fA)#(mU)#(fA) | 6657.61 | 6657.59 |
| **G33** | 3-(4-pyridyl)propargyl-P(mU)#(fU)#(mA)(fA)(mU)(fC)(mU)(fC)(mU)(fU)(mU)(fA)(mC)#(fU)#(mG)#(fA)#(mU)#(fA)#(mU)#(fA) | 6657.61 | 6657.50 |
| **G34** | 3-(2-thienyl)propargyl-P(mU)#(fU)#(mA)(fA)(mU)(fC)(mU)(fC)(mU)(fU)(mU)(fA)(mC)#(fU)#(mG)#(fA)#(mU)#(fA)#(mU)#(fA) | 6662.65 | 6662.04 |
| **G35** | 3-aminopropyl-P(mU)#(fU)#(mA)(fA)(mU)(fC)(mU)(fC)(mU)(fU)(mU)(fA)(mC)#(fU)#(mG)#(fA)#(mU)#(fA)#(mU)#(fA) | 6599.57 | 6599.42 |
| **G36** | 3-acetamidopropyl-P(mU)#(fU)#(mA)(fA)(mU)(fC)(mU)(fC)(mU)(fU)(mU)(fA)(mC)#(fU)#(mG)#(fA)#(mU)#(fA)#(mU)#(fA) | 6641.61 | 6640.84 |
| **G37** | PS(mU)#(fU)#(mA)(fA)(mU)(fC)(mU)(fC)(mU)(fU)(mU)(fA)(mC)#(fU)#(mG)#(fA)#(mU)#(fA)#(mU)#(fA) | 6636.51 | 6636.02 |
| **G38** | phenylpropargyl-PS(mU)#(fU)#(mA)(fA)(mU)(fC)(mU)(fC)(mU)(fU)(mU)(fA)(mC)#(fU)#(mG)#(fA)#(mU)#(fA)#(mU)#(fA) | 6750.65 | 6750.60 |
| **G39** | MsPA(mU)#(fU)#(mA)(fA)(mU)(fC)(mU)(fC)(mU)(fU)(mU)(fA)(mC)#(fU)#(mG)#(fA)#(mU)#(fA)#(mU)#(fA) | 6697.54 | 6696.77 |
| **G40** | P(mU)#(fA)#(mU)(fC)(mG)(fG)(mG)(fA)(mA)(fG)(mC)(fU)(mU)#(fU)#(mG)#(fU)#(mC)#(fA)#(mG)#(fA) | 6792.60 | 6792.12 |
| **G41** | methyl-(mU)#(fA)#(mU)(fC)(mG)(fG)(mG)(fA)(mA)(fG)(mC)(fU)(mU)#(fU)#(mG)#(fU)#(mC)#(fA)#(mG)#(fA) | 6806.62 | 6806.36 |
| **G42** | phenylpropargyl-P(mU)#(fA)#(mU)(fC)(mG)(fG)(mG)(fA)(mA)(fG)(mC)(fU)(mU)#(fU)#(mG)#(fU)#(mC)#(fA)#(mG)#(fA) | 6906.73 | 6906.55 |
| **G43** | (mU)#(fU)#(mA)(fA)(mU)(fC)(mU)(fC)(mU)(fU)(mU)(fA)(mC)#(fU)#(mG)#(fA)#(mU)#(fA)#(mU)#(fA) | 6540.46 | 6539.41 |
| **P1** | (fC)#(mA)#(fG)(mU)(fA)(mA)(fA)(mG)(fA)(mG)(fA)(mU)(fU)#(mA)#(fA)Chol | 5765.34 | 5765.29 |
| **P2** | (fC)#(mA)#(fG)(mU)(fA)(mA)(fA)(mG)(fA)(mG)(fA)(mU)(fU)#(mA)#(fA) | 5009.37 | 5008.62 |
| **P3** | (fC)#(mA)#(fA)(mA)(fG)(mC)(fU)(mU)(fC)(mC)(fC)(mG)(fA)#(mU)#(fA) | 4897.26 | 4896.55 |
| **ON1** | P(mU)(fU)(mA)(fA)(mU)(fC)(mU)(fC)(mU)(fU)(mU)(fA)(mC)(fU)(mG)(fA)(mU)(fA)(mU)(fA) | 6475.84 | 6474.86 |

| | | | |
|---|---|---|---|
| **ON2** | (mU)(fU)(mA)(fA)(mU)(fC)(mU)(fC)(mU)(fU)(mU)(fA)(mC)(fU)(mG)(fA)(mU)(fA)(mU)(fA) | 6,395.86 | 6395.58 |
| **ON3** | VP(mU)(fU)(mA)(fA)(mU)(fC)(mU)(fC)(mU)(fU)(mU)(fA)(mC)(fU)(mG)(fA)(mU)(fA)(mU)(fA) | 6,470.84 | 6470.64 |
| **ON4** | methyl-P(mU)(fU)(mA)(fA)(mU)(fC)(mU)(fC)(mU)(fU)(mU)(fA)(mC)(fU)(mG)(fA)(mU)(fA)(mU)(fA) | 6489.87 | 6489.78 |
| **ON5** | phenylpropargyl-P(mU)(fU)(mA)(fA)(mU)(fC)(mU)(fC)(mU)(fU)(mU)(fA)(mC)(fU)(mG)(fA)(mU)(fA)(mU)(fA) | 6589.99 | 6589.39 |
| **ON6** | PS(mU)(fU)(mA)(fA)(mU)(fC)(mU)(fC)(mU)(fU)(mU)(fA)(mC)(fU)(mG)(fA)(mU)(fA)(mU)(fA) | 6491.91 | 6491.47 |
| **ON7** | phenylpropargyl-PS(mU)(fU)(mA)(fA)(mU)(fC)(mU)(fC)(mU)(fU)(mU)(fA)(mC)(fU)(mG)(fA)(mU)(fA)(mU)(fA) | 6606.95 | 6606.43 |
| **ON8** | MsPA(mU)(fU)(mA)(fA)(mU)(fC)(mU)(fC)(mU)(fU)(mU)(fA)(mC)(fU)(mG)(fA)(mU)(fA)(mU)(fA) | 6552.94 | 6552.39 |
| **ON9** | (dT)(dT)(dT)(dT)(dT)(dT)(dT)(dT) | 2371.59 | 2371.24 |
| **ON10** | phenylpropargyl-P(mU)#(fG)#(mG)(fA)(fG)(fU)(mG)(fU)(mG)(fA)(mC)(fA)(mA)(fU)(mG)(fG)(mU)(mG)(mU)#(mU)#(fU) | 7203.62 | 7203.13 |
| **ON11** | 3-phenylpropyl-P(mU)#(fG)#(mG)(fA)(fG)(fU)(mG)(fU)(mG)(fA)(mC)(fA)(mA)(fU)(mG)(fG)(mU)(mG)(mU)#(mU)#(fU) | 7207.65 | 7207.34 |
| **ON12** | methyl-P(mU)#(fG)#(mG)(fA)(fG)(fU)(mG)(fU)(mG)(fA)(mC)(fA)(mA)(fU)(mG)(fG)(mU)(mG)(mU)#(mU)#(fU) | 7103.50 | 7103.69 |
| **ON13** | P(mU)#(fG)#(mG)(fA)(fG)(fU)(mG)(fU)(mG)(fA)(mC)(fA)(mA)(fU)(mG)(fG)(mU)(mG)(mU)#(mU)#(fU) | 7089.47 | 7089.21 |

Chemical modifications are designated as follows. "#": phosphorothioate linkage; "f": 2′-deoxy-2′-fluoro; "m": 2′-*O*-methyl; "d": 2′-deoxy; "P": 5′-phosphate; "PS": 5′-phosphorothioate; "VP": 5′-(*E*)-vinylphosphonate; "MsPA"; 5′-mesylphosphoramidate; "Chol": 3′-cholesterol conjugate (ChemGenes). Any substituents are named using standard chemical nomenclature.
